# Mouse retinal specializations reflect knowledge of natural environment statistics

**DOI:** 10.1101/2020.12.08.416172

**Authors:** Yongrong Qiu, Zhijian Zhao, David Klindt, Magdalena Kautzky, Klaudia P. Szatko, Frank Schaeffel, Katharina Rifai, Katrin Franke, Laura Busse, Thomas Euler

**Author notes:** Lead contact: Thomas Euler.

## Abstract

Pressures for survival drive sensory circuit adaption to a species’ habitat, making it essential to statistically characterise natural scenes. Mice, a prominent visual system model, are dichromatic with enhanced sensitivity to green and UV. Their visual environment, however, is rarely considered. Here, we built a UV-green camera to record footage from mouse habitats. We found chromatic contrast to greatly diverge in the upper but not the lower visual field, an environmental difference that may underlie the species’ superior colour discrimination in the upper visual field. Moreover, training an autoencoder on upper but not lower visual field scenes was sufficient for the emergence of colour-opponent filters. Furthermore, the upper visual field was biased towards dark UV contrasts, paralleled by more light-offset-sensitive cells in the ventral retina. Finally, footage recorded at twilight suggests that UV promotes aerial predator detection. Our findings support that natural scene statistics shaped early visual processing in evolution.

**Lead contact:** Further information and requests for resources and reagents should be directed to and will be fulfilled by the Lead Contact, Thomas Euler

## Introduction

During evolution, the structure and function of neural circuits have been shaped to improve the species’ chances to survive and procreate in their specific natural environments. Such adaptations have long been described and discussed in the visual system (Attneave, 1954; Barlow, 1961; Simoncelli and Olshausen, 2001). Some of these adaptations are already in eye placement: For example, predators, such as cats, usually have frontally placed eyes, maximizing the binocular field to improve stereovision. Prey animals, such as mice, have laterally placed eyes, expanding the field-of-view (FOV) to detect threats as reliably as possible (reviewed in Baden et al., 2020).

Given such specific adaptations, characterising the properties of natural visual environments is crucial for advancing our understanding of the structure and function of the visual system (Masland and Martin, 2007), in particular for species whose visual systems are different from ours, such as mice. Yet, studying the visual system and behaviour of mice in the context of their natural environment is only starting (Datta et al., 2019; Hasson et al., 2020; reviewed in Krakauer et al., 2017), despite the fact that mice have become a prominent model system for vision research in the past decade (reviewed in Huberman and Niell, 2011; Seabrook et al., 2017). The importance of considering ethologically relevant stimuli and behaviours for studying mouse vision is highlighted in recent work, showing, for example, superior spatial frequency tuning for V1 neurons when probed with ecologically inspired visual stimuli instead of drifting gratings (Dyballa et al., 2018), which might potentially underlie accurate visually-driven approach performance during prey capture (Hoy et al., 2016).

The relationship between properties of natural scenes and principles of visual coding has been powerfully addressed using computational modeling (reviewed in Turner et al., 2019). A classic example is the finding that simple cell-like receptive fields (RFs) emerge in models that learn a sparse code for natural images (Olshausen and Field, 1996). Along these lines, Ocko and coworkers (2018) recently showed that a convolutional autoencoder (CAE; Ballard, 1987; Hinton and Salakhutdinov, 2006) trained to reconstruct pink (1/*f*) noise learned center-surround spatial filters reminiscent of the RFs of different retinal ganglion cell (RGC) types. In addition, imposing constraints on encoding models to drive optimal performance in specific natural vision tasks, has proven fruitful to generate principled hypotheses about the underlying neural or perceptual mechanisms (Burge and Jaini, 2017; Geisler et al., 2009). Such a task-driven framework has been fueled in the last years by advances in deep convolutional neural networks (DNNs) based on discriminative learning from large databases of natural images (reviewed in LeCun et al., 2015; Turner et al., 2019). Interestingly, such layered networks learn features that share representational similarities with neuronal activity in several processing stages along the visual hierarchy (Yamins and DiCarlo, 2016).

In natural scenes, colour is an ethologically highly relevant visual feature, and depending on which colours an animal can see, this will influence its ability to forage and hunt, select mates, and avoid predators. Indeed, beyond the absence of a fovea in mice, a major difference compared to primates is that mice are dichromats and see UV light. Next to a medium (M) wavelength-sensitive opsin peaking at 510 nm (green), mice have a short (S) wavelength-sensitive opsin peaking at 360 nm in the UV (Jacobs et al., 2004; Röhlich et al., 1994). Due to co-expression of the S-opsin in ventral M-cones (Baden et al., 2013; Szél et al., 1992) and a peak in S-cone density in the ventral periphery (Nadal-Nicolás et al., 2020), the mouse retina is subdivided into a more green-sensitive dorsal and a strongly UV-sensitive ventral half. Notably, ventral cones are tuned to signal dark contrasts (Baden et al., 2013) and, hence, may support the detection of dark shapes against the sky. Such dorso-ventral regionalization of the mouse retina suggests a functional specialization, according to which the ventral retina “monitors” the overhead space to detect predators (Wallace et al., 2013; Zhang et al., 2012), whereas the dorsal retina supports foraging and hunting for food (Hoy et al., 2019; Shang et al., 2019). An important consequence of this regionalization is that any statistical analysis of the mouse’s visual environment — like quantifying spectral content of natural scenes — should consider the upper and lower visual field separately. However, recording the chromatic information available to mice under natural conditions is challenging, since standard cameras fail to capture UV light.

To analyze the spectral properties of the environment of mice, we have therefore built a hand-held camera that is mounted on a gimbal for image stabilization and covers the spectral bands – UV and green – relevant for mice. We used this camera to record footage near mouse tracks in the field, capturing various outside scenes at different times of the day – providing a resource of natural scenes for vision research in mice. Focusing on spectral information, we found that the contrast in the two spectral channels greatly diverged in the upper but not in the lower visual field, paralleling the superior chromatic opponency in the ventral retina (Szatko et al., 2020) and behavioural colour discrimination in the upper visual field (Denman et al., 2018). Notably, our analysis of the footage suggests that the mouse’s UV sensitivity may help detecting aerial predators also at dusk and dawn. Finally, computational modelling predicts that colour-opponent filters are more likely to emerge in unsupervised models trained with images from the upper visual field than from the lower visual field. Together, this lends further support to the idea that retinal circuits have evolved to process natural scene statistics in a species-specific manner (reviewed in Baden et al., 2020).

## Results

### A camera for recording visual scenes from the mouse’s perspective

The goal of this study was to capture the visual environment of mice mimicking key aspects of mouse vision. Specifically, we focused on three main aspects: (i) the perspective from only a few centimeters above the ground, (ii) the large FOV that for each eye approaches ~180° (Sterratt et al., 2013), and (iii) the spectral sensitivities of the mouse photoreceptors, peaking at 360 and 510 nm for UV-cones and rods/M-cones, respectively.

To this end, we developed a “mouse-camera” that simultaneously captures movies in the UV and green spectral bands (Fig. 1; for components, see Table 1; for camera settings, see Table 2; for additional details, see Methods). The two spectral channels were simultaneously recorded on flash memory by two single-board Raspberry Pi microcomputers attached to the camera modules. As an objective, we used a fisheye lens with a FOV of 180° (Fig. 1a-e). The camera was mounted on an active gimbal for stabilization, such that it could be moved close to the ground (Fig. 1c-e).

**Figure 1.**
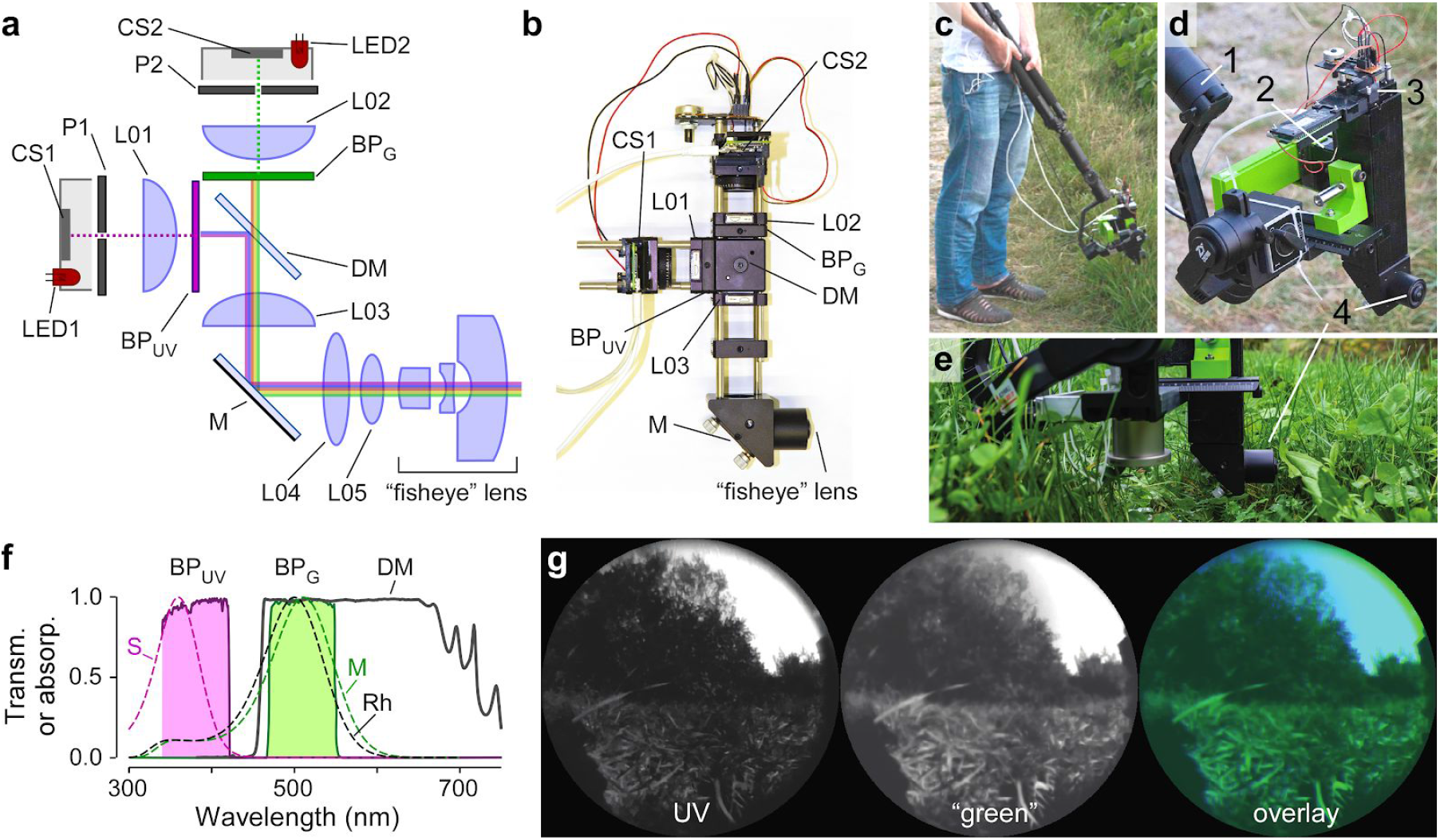
Mouse-camera module. **a**, Schematic drawing of mouse-camera with two spectral channels (UV, green). CS, camera sensor; P, pinhole; L01-05, lenses; BP_UV_, UV bandpass filter (350-419 nm); M, silver mirror; LED, light-emitting diode; BP_G_ green bandpass filter (470-550 nm); DM, dichroic mirror (>90% reflection: 350-442 nm; >90%, transmission: 460-650 nm). **b**, Picture of the assembled camera module. **c-e**, Pictures of assembled camera module, with gimbal (1), UV camera (2), green camera (3), and fisheye lens (4). **f**, Normalized transmission spectra of DM, BP_UV_ and BP_G_, with normalized absorption spectra of mouse cone opsins (S, M) and rhodopsin (Rh) overlaid (Franke et al., 2019). **g**, Movie frame with UV (left), green (centre) channel, and overlay (right).

**Table 1.**
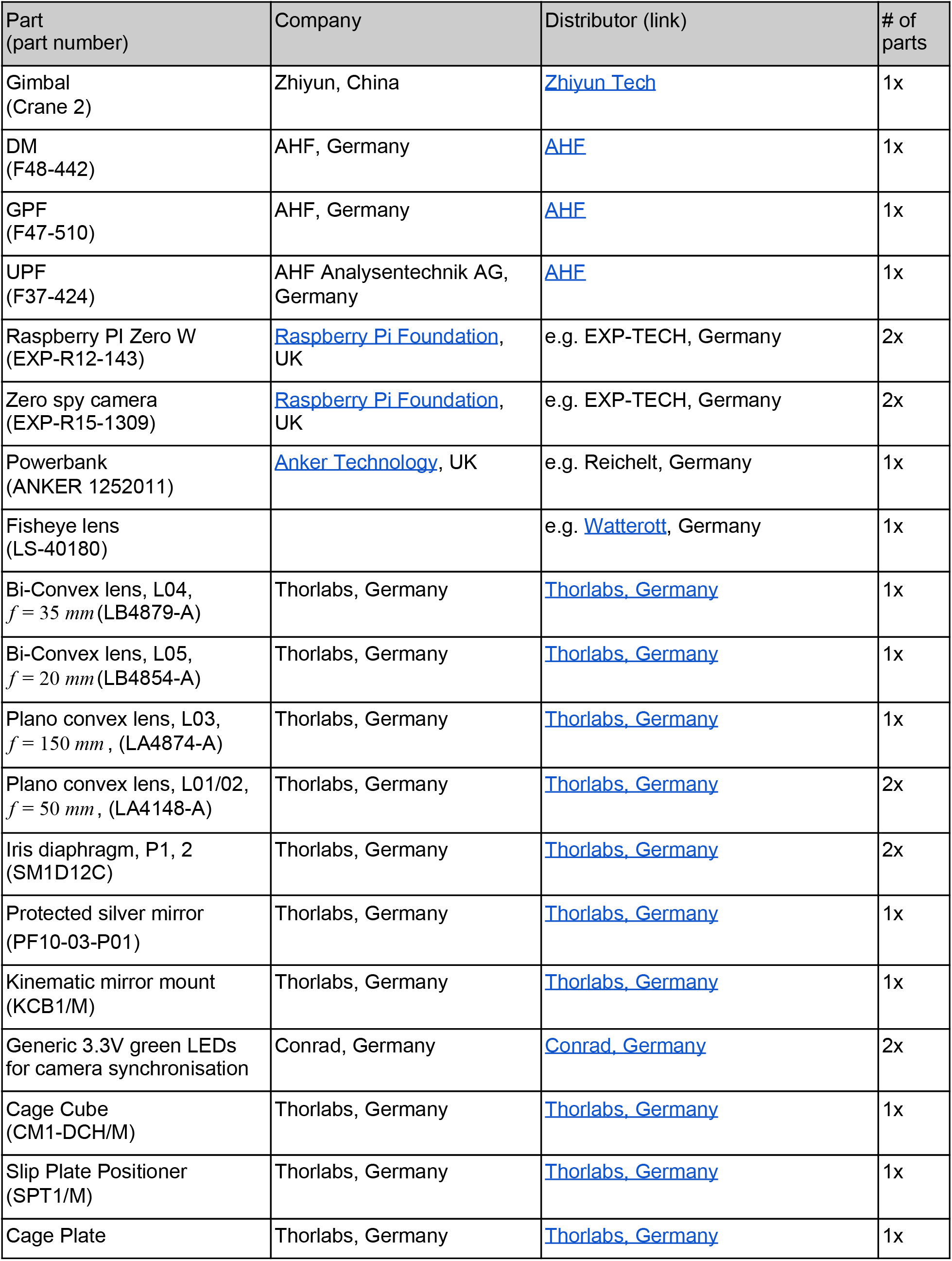
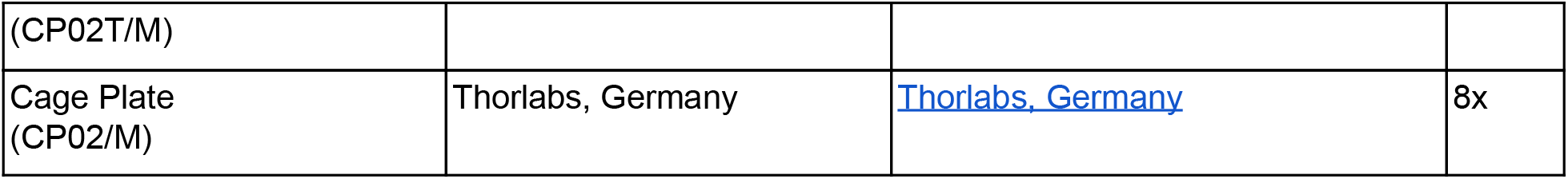
Camera parts.

**Table 2.**
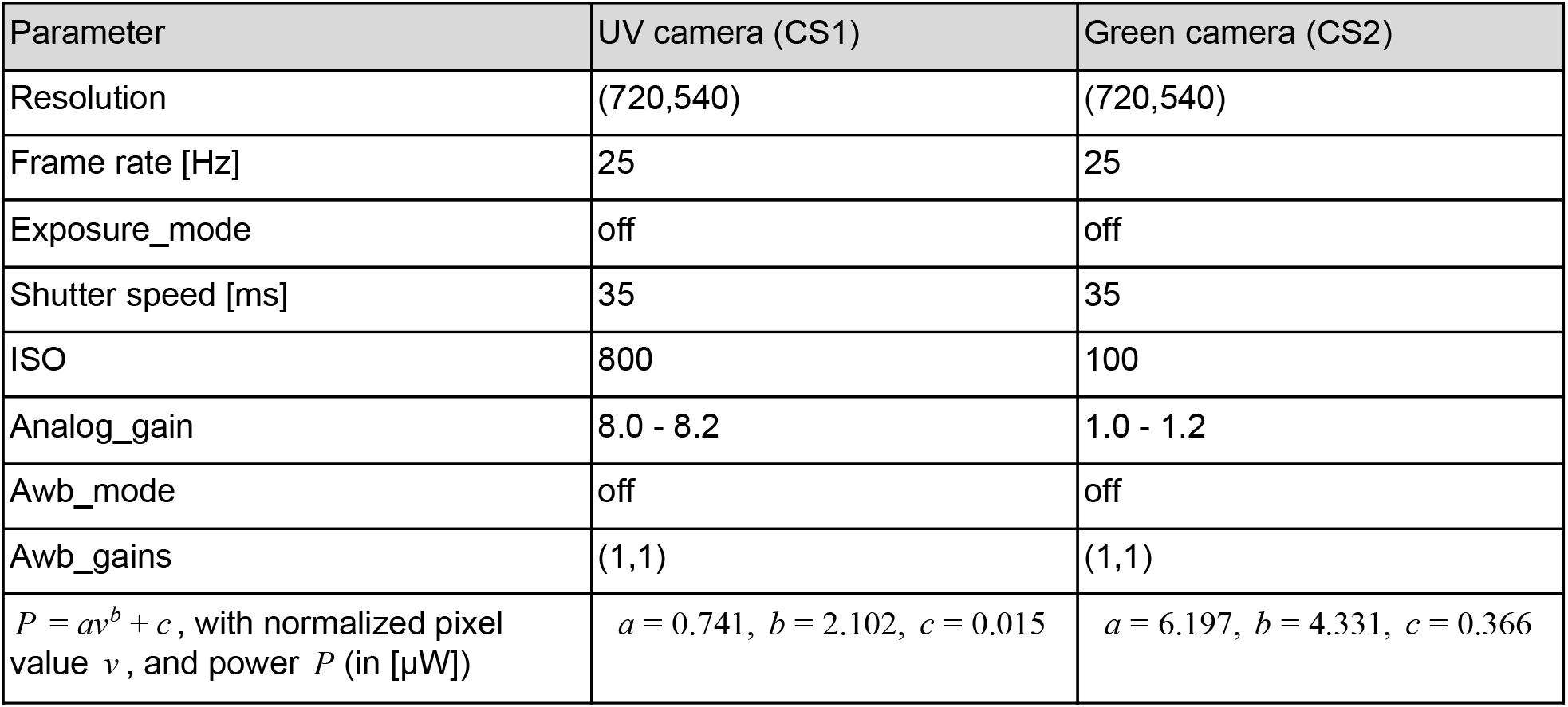
Camera settings.

| Parameter | UV camera (CS1) | Green camera (CS2) |
| --- | --- | --- |
| Resolution | (720,540) | (720,540) |
| Frame rate [Hz] | 25 | 25 |
| Exposure_mode | off | off |
| Shutter speed [ms] | 35 | 35 |
| ISO | 800 | 100 |
| Analog_gain | 8.0 - 8.2 | 1.0 - 1.2 |
| Awb_mode | off | off |
| Awb_gains | (1,1) | (1,1) |
| $P = av^b + c$, with normalized pixel value $v$, and power $P$ (in [μW]) | $a = 0.741$, $b = 2.102$, $c = 0.015$ | $a = 6.197$, $b = 4.331$, $c = 0.366$ |

Based on the assumption that eye movements in mice serve mainly to stabilize the retinal image (Land, 2019; Meyer et al., 2018, 2020; Michaiel et al., 2020) and are typically coupled to head movements (Meyer et al., 2020; Michaiel et al., 2020), we restricted our recordings to a view with the horizon oriented parallel to ground and positioned around the middle elevation of the camera image. This way, the lower half of the camera image captured information from the lower visual field of mice, while the upper half captured information from the upper visual field.

The Raspberry Pi camera modules we used are consumer products (for specifications, see Table 3) and, hence, are optimized for taking colour pictures that look natural to a human observer. At the hardware side, this is achieved by a checkerboard-like pattern of thin RGB filters (Bayer filter) coated directly on the chip surface (for spectra, see resources listed in Table 3). While the Bayer filter’s green component matches the spectral sensitivity of the mouse rods and M-cones (λ_*Peak*_ ≈ 510 *nm*) (Jacobs et al., 2004, 2007), the Bayer filter largely blocks UV light. To address this issue, we mechanically removed the Bayer filter layer in case of the UV channel camera chip (Wilkes et al., 2016).

**Table 3.**
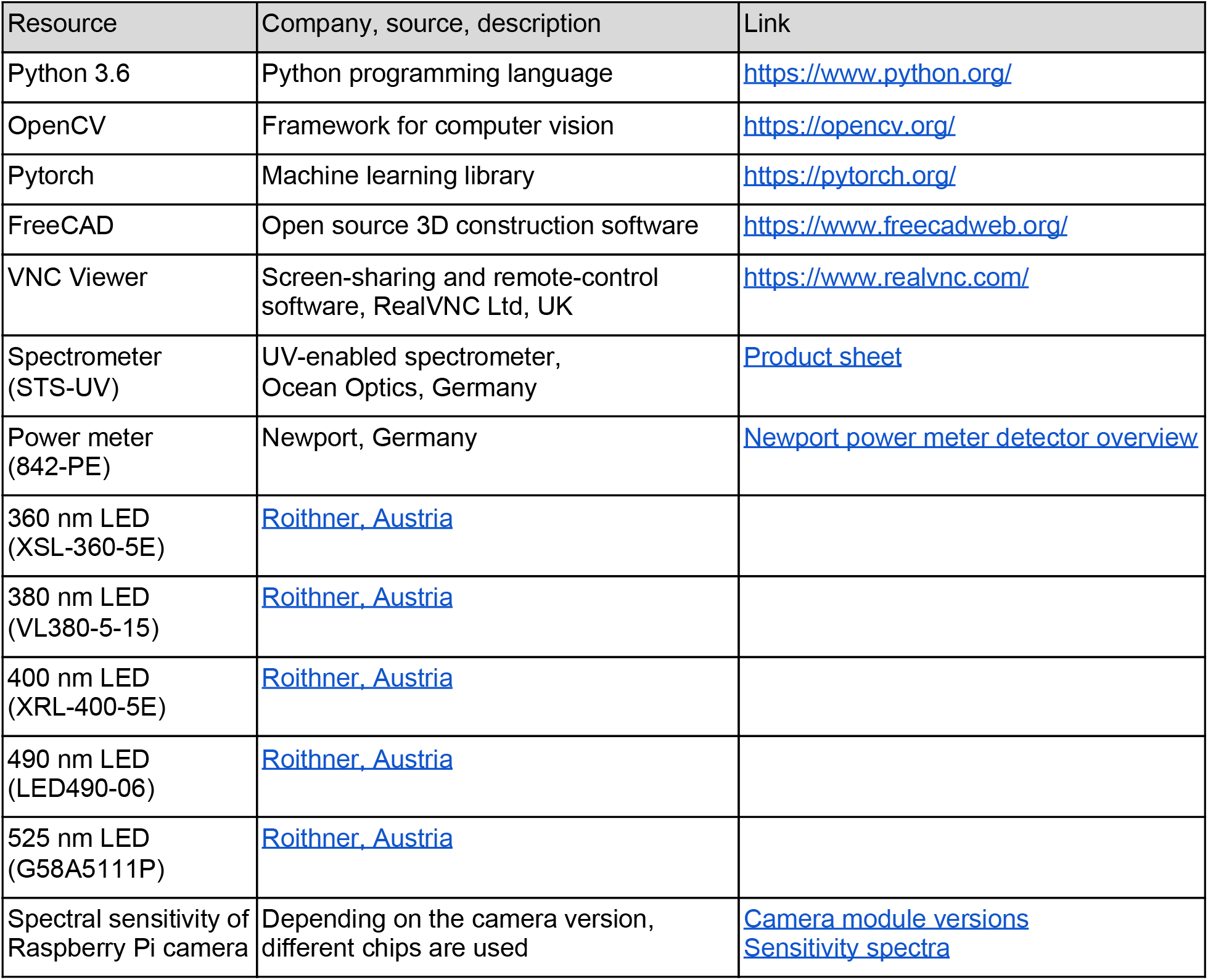
Resources (except camera parts).

| Resource | Company, source, description | Link |
| --- | --- | --- |
| Python 3.6 | Python programming language | https://www.python.org/ |
| OpenCV | Framework for computer vision | https://opencv.org/ |
| Pytorch | Machine learning library | https://pytorch.org/ |
| FreeCAD | Open source 3D construction software | https://www.freecadweb.org/ |
| VNC Viewer | Screen-sharing and remote-control software, RealVNC Ltd, UK | https://www.realvnc.com/ |
| Spectrometer (STS-UV) | UV-enabled spectrometer, Ocean Optics, Germany | Product sheet |
| Power meter (842-PE) | Newport, Germany | Newport power meter detector overview |
| 360 nm LED (XSL-360-5E) | Roithner, Austria |  |
| 380 nm LED (VL380-5-15) | Roithner, Austria |  |
| 400 nm LED (XRL-400-5E) | Roithner, Austria |  |
| 490 nm LED (LED490-06) | Roithner, Austria |  |
| 525 nm LED (G58A5111P) | Roithner, Austria |  |
| Spectral sensitivity of Raspberry Pi camera | Depending on the camera version, different chips are used | Camera module versions<br>Sensitivity spectra |

The effective spectral sensitivity of each camera channel is determined by the combination of bare chip sensitivity, Bayer filter (in case of the green channel), and a spectral bandpass filter (Fig. 1f). In addition, the cameras automatically adjust the image intensities to match gamma curves typically found in consumer displays. Since we do not know the chip’s exact sensitivity curve, we measured the sensitivity of the two camera channels using LEDs of defined spectrum and power (Suppl. Figure S1a; Methods). For each channel, we mapped image intensities to power meter readouts of UV or green LEDs, and fitted the data using an inverse gamma correction function (Suppl. Fig. S1b). We then used these fits to convert pixel values into normalized intensities (source vs. intensity-corrected images; see Suppl. Fig. S1c).

To verify this intensity correction, we recorded outside scenes using a scanning spectrometer (Baden et al., 2013), which provides still images with each pixel containing the spectrum from 300 to 660 nm (see Methods). By convolving each pixel’s spectrum with the mouse cone opsin absorption curves, we generated mouse-view corrected versions of the spectrometer images (Suppl. Fig. S1d). For representative scene elements, such as grass and trees, we found that the normalized intensities in the corrected camera images matched well those in the spectrometer images for both the UV and green channel (Suppl. Fig. S1e).

Note that the intensity correction reached its limits for extensive sky regions, because the camera chip could saturate due to its limited dynamic range, when the spectrometer did not. We considered this potential issue by excluding oversaturated images (see Methods). Note also that we used the available range for both colour channels by mapping the intensities to pixel values (see Methods), although in the natural environment, the absolute intensity of UV is known to be lower than that of longer wavelengths (Hut et al., 2000; see Discussion). After intensity correction, the movies from the two channels were temporally synchronized and spatially overlayed (see Methods), resulting in a single UV-green movie (Suppl. Video V1).

### Movie recordings and first-order statistics

Next, we used the camera to record footage of representative natural scenes outside in the field in places with traces of mouse activity (Fig. 2a). Most scenes were recorded in summer and spring during the day, and a few scenes at dusk or dawn (*cf*. Fig. 5; see Table 4).

**Figure 2.**
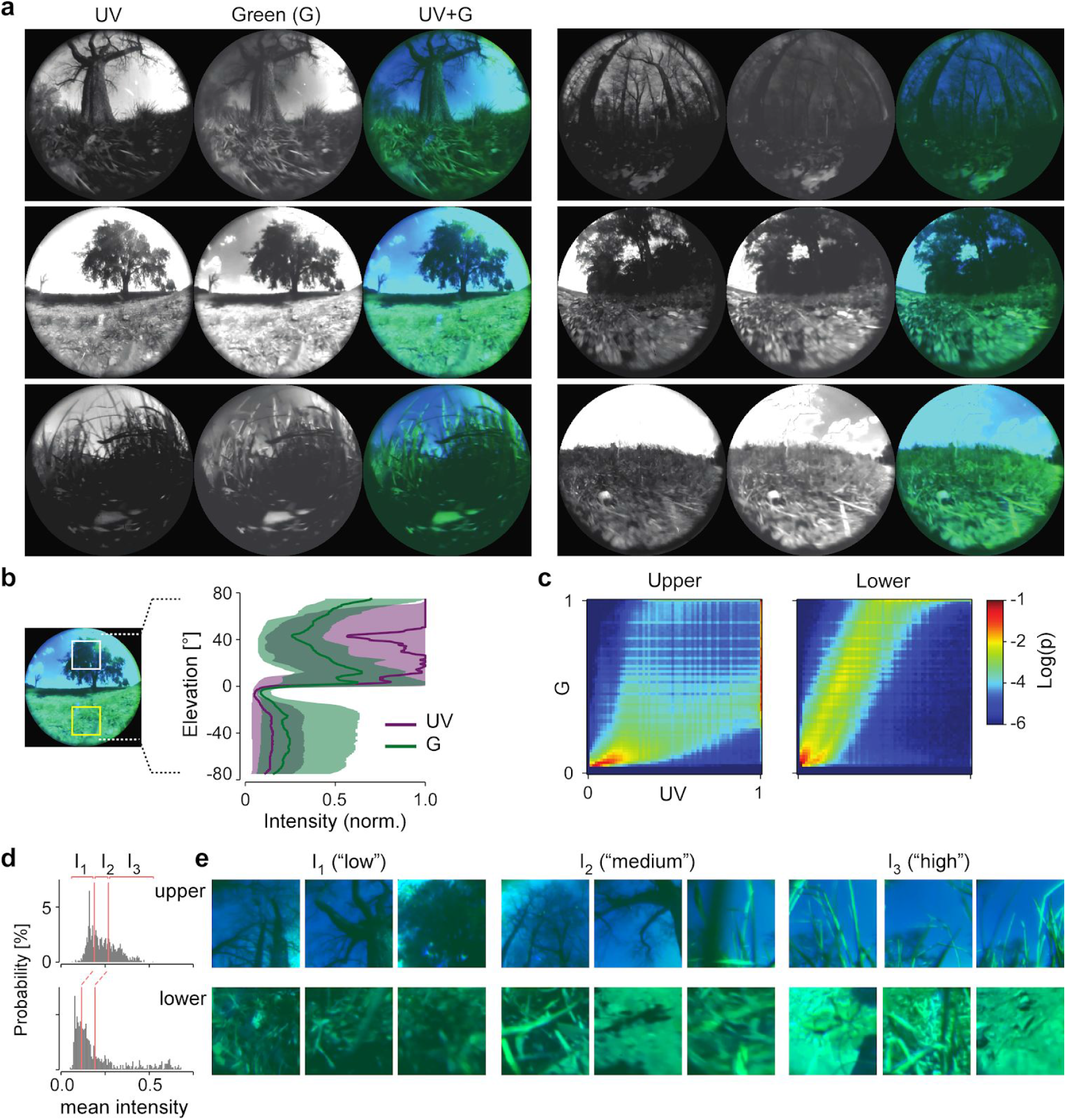
Example scenes and intensity distribution. **a**, Example frames (UV, green, and overlay) from movies of different scenes recorded outside in the field (near Waldhäuser Ost, Tübingen, Germany; 48°33’02.4”N 9°03’01.2”E). **b**, Normalized intensity for green (G) and UV as a function of elevation across the n=1,936 frames of one example movie (median w/ 25^th^ and 75^th^ percentile). **c**, 2D histograms of intensities of the same movie as in (b), visualizing the G-UV intensity distribution for an image cut-out (crop) in the upper (left) and lower (right) visual field (for region placement, see yellow boxes in (b, left)). **d,** Distribution of mean intensities for 1,500 image crops from upper and lower visual field, selected randomly from 15 movies, and division into 3 intensity classes (percentiles I_1-3_). **e**, Example images from the three mean intensity classes.

**Table 4.**
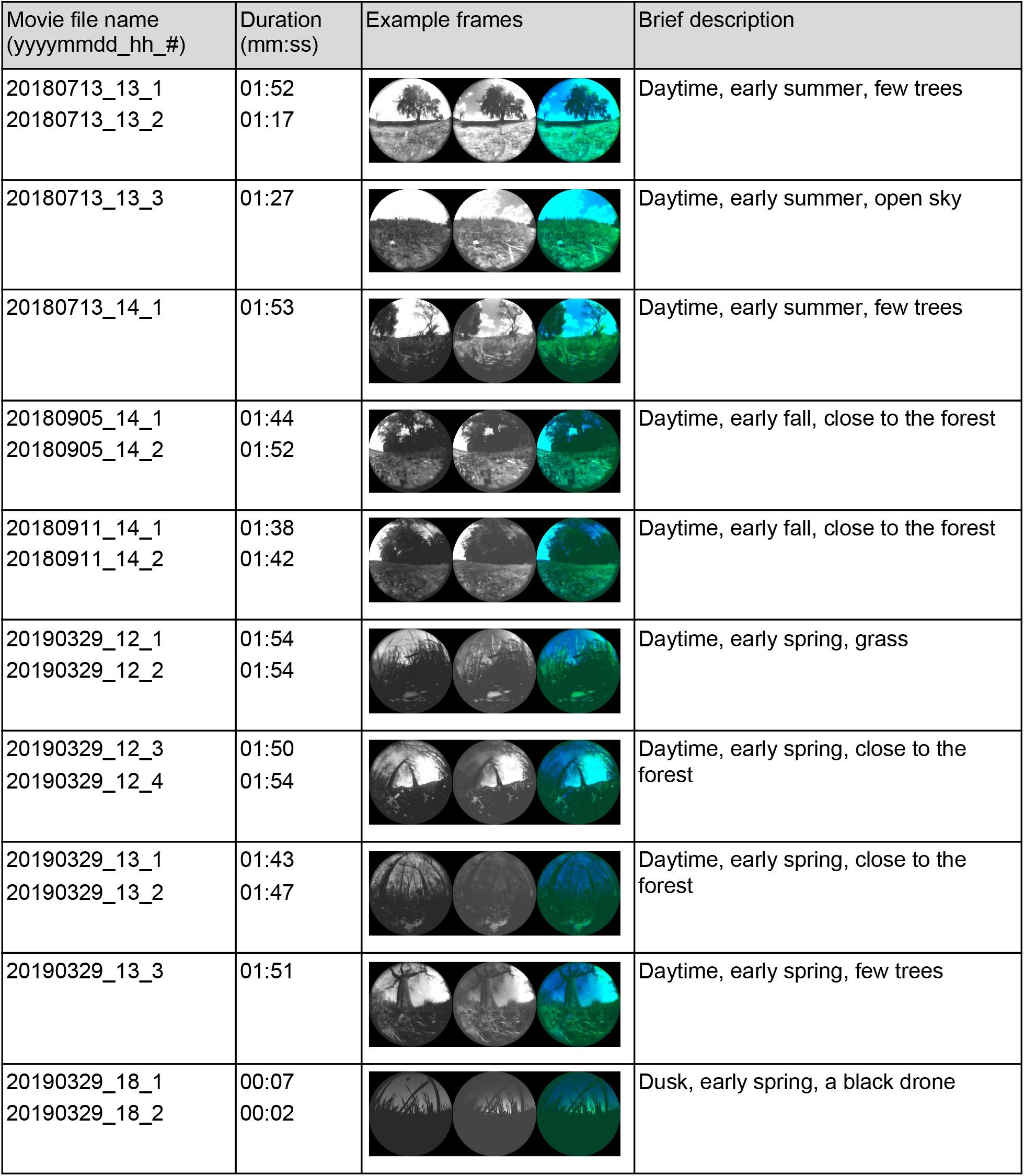
Natural movies.

| Movie file name<br>(yyyymmdd_hh_#) | Duration<br>(mm:ss) | Example frames | Brief description |
| --- | --- | --- | --- |
| 20180713_13_1<br>20180713_13_2 | 01:52<br>01:17 |  | Daytime, early summer, few trees |
| 20180713_13_3 | 01:27 |  | Daytime, early summer, open sky |
| 20180713_14_1 | 01:53 |  | Daytime, early summer, few trees |
| 20180905_14_1<br>20180905_14_2 | 01:44<br>01:52 |  | Daytime, early fall, close to the forest |
| 20180911_14_1<br>20180911_14_2 | 01:38<br>01:42 |  | Daytime, early fall, close to the forest |
| 20190329_12_1<br>20190329_12_2 | 01:54<br>01:54 |  | Daytime, early spring, grass |
| 20190329_12_3<br>20190329_12_4 | 01:50<br>01:54 |  | Daytime, early spring, close to the forest |
| 20190329_13_1<br>20190329_13_2 | 01:43<br>01:47 |  | Daytime, early spring, close to the forest |
| 20190329_13_3 | 01:51 |  | Daytime, early spring, few trees |
| 20190329_18_1<br>20190329_18_2 | 00:07<br>00:02 |  | Dusk, early spring, a black drone |

We first focused on first-order statistics, i.e. brightness, and explored the normalized intensity distribution in each channel as a function of elevation (Fig. 2b). Examining crops from an example movie, we found that the relative intensities in both channels were usually higher in the upper compared to the lower visual field (Fig. 2b, right). Interestingly, the two chromatic channels were less correlated in image crops from the upper than the lower visual field (Fig. 2c). We therefore quantified the linear correlation between the two channels using principal component analysis (PCA; Buchsbaum and Gottschalk, 1983) and found a 2-5 times higher variance along the colour-opponent axis in images from the upper vs. the lower visual field (see colour opponency index *CO_i_* in Suppl. Fig. S2). This indicates a higher variability in chromatic intensity differences, i.e. contrasts, above the horizon (see also next section).

Performing these analyses systematically across all recorded movies comes with the challenge that, depending on the time of the day, the weather, and the scene content, the brightness of the recorded scenes could vary tremendously – for instance when comparing a scene with clear sky and a single tree (Fig. 2a, left-centre) with a scene recorded close to the forest (right-top). We therefore split the footage crop-wise into three groups based on the mean intensity (Fig. 2d,e; see Methods) and performed all subsequent analyses for each of these brightness categories.

### Chromatic contrast is higher in the upper vs. the lower visual field

Retinal output to the brain is also driven by second-order statistics, such as differences in brightness, i.e. contrast (Frazor and Geisler, 2006; Mante et al., 2005; Rieke and Rudd, 2009). If distributions of contrast differ between colour channels, this difference can be interpreted as chromatic contrast. Therefore, we next compared the contrast distribution in the chromatic channels in our natural scenes above and below the horizon and across the three mean intensity groups (Fig. 3).

**Figure 3.**
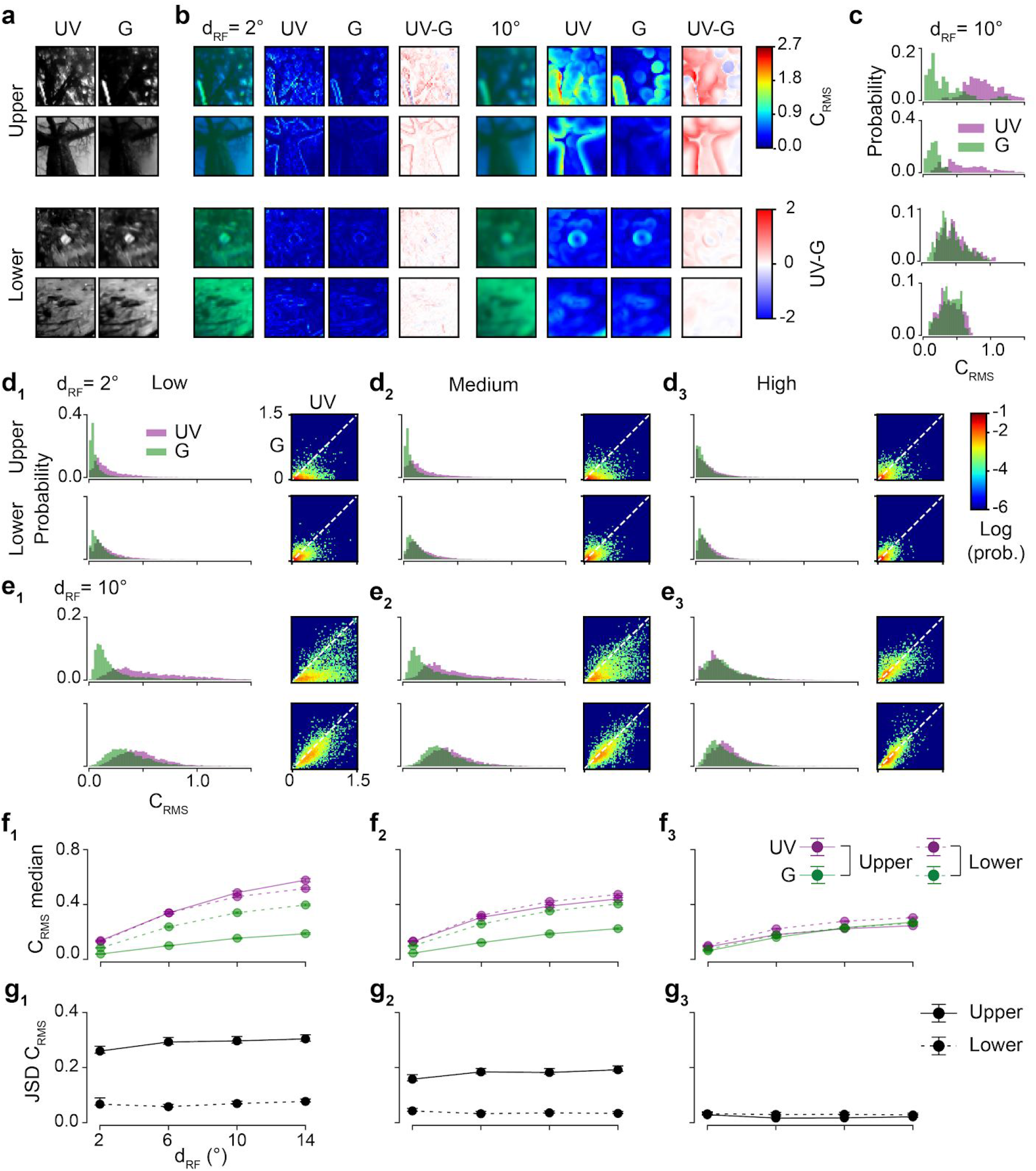
Chromatic contrast is higher in image crops from the upper visual field. **a,** Examples for image crops from upper (from I_1_ and I_2_) and lower (from I_1_ and I_3_) visual field. **b**, images from (a) filtered with different receptive field (RF) diameters, d_RF_, with each of the columns showing (from left): UV-green overlay, UV and green channel maps visualizing RMS contrast (C_RMS_), and difference between maps (UV-G). **c**, Distributions of C_RMS_ for UV and green for d_RF_=10°. **d,e**, 1D (left) and 2D histograms for C_RMS_ of all n=1,500 image crops from randomly picked frames (out of n=15 movies) for the three intensity groups (from Fig. 2d,e) and d _RF_=2° (d) and d_RF_=10° (e). **f**, Median C _RMS_ as function of RF size for UV and green channels in the upper (solid lines) and lower (dashed) visual field, for the three intensity groups. **g**, Like (f) but with median Jensen-Shannon divergence (JSD) between C_RMS_ distributions of the two chromatic channels as function of d_RF_. Error bars in (f,g) represent 2.5 and 97.5 percentiles with bootstrapping (see Methods).

We first focused on root mean square contrast (*C_RMS_*), commonly used in psychophysics for describing contrast in complex natural scenes (Bex and Makous, 2002; Mante et al., 2005). We extracted circular image patches (kernels) of various diameters (*d_RF_*, 2° to 14°), a range that includes the RF sizes of mouse RGC types (Baden et al., 2016), and computed *C_RMS_* as the standard deviation of the normalized pixel intensities divided by mean intensity. To quantify differences between *C_RMS_* distributions, we used a two-sided permutation test as well as the Jensen-Shannon divergence (*JSD*), which measures the general similarity between probability distributions, with *JSD* = 0 indicating identical distributions.

Our analysis of the *C_RMS_* distributions across 1,500 crops revealed three key features of the recorded mouse natural scenes (Fig. 3): First, as illustrated by two representative examples (Fig. 3a-c; upper two vs. lower two rows; from Suppl. Video V1,2), *C_RMS_* was higher in the upper compared to the lower visual field (for *C_RMS_* distributions, medians and JSD, see Fig. 3d-g). This difference was much more pronounced in the UV than the green channel and in the low and medium mean intensity groups (compare columns in Fig. 3d-g). Second, for the tested kernel sizes, *C_RMS_* increased with kernel diameter (see examples in Fig. 3b; compare Fig. 3d vs. e; for median and JSD, see Fig. 3f,g), consistent with the dominance of low-spatial frequencies in natural scenes (Burton and Moorhead, 1987; Field, 1987; Ruderman and Bialek, 1994). Third, *C_RMS_* distributions of the UV and green channel differed more strongly in the upper compared to the lower visual field (see example in Fig. 3c; for median and JSD, see Fig. 3f,g). These differences in *C_RMS_* distribution between chromatic channels and upper vs. lower visual field were significant for all kernel diameters except in the high mean intensity group (for all comparisons, see Tables 5, 6).

**Table 5.** Statistics of C_RMS_ between chromatic channels and upper vs. lower visual field as a function of kernel diameter. Two-sided permutation test with 10,000 repeats. White, light orange and dark orange indicating p < 0.0001, 0.0001 ≤ p < 0.05 and p ≤ 0.05

| Group | $d_{\text{RF}}(^{\circ})$ | Upper UV<br>vs.<br>Upper G | Upper UV<br>vs.<br>Lower UV | Upper UV<br>vs.<br>Lower G | Upper G<br>vs.<br>Lower UV | Upper G<br>vs.<br>Lower G | Lower UV<br>vs.<br>Lower G |
| --- | --- | --- | --- | --- | --- | --- | --- |
| $I_1$ | 2 | <0.0001 | <0.0001 | <0.0001 | <0.0001 | <0.0001 | <0.0001 |
|  | 6 | <0.0001 | <0.0001 | <0.0001 | <0.0001 | <0.0001 | <0.0001 |
|  | 10 | <0.0001 | <0.0001 | <0.0001 | <0.0001 | <0.0001 | <0.0001 |
|  | 14 | <0.0001 | <0.0001 | <0.0001 | <0.0001 | <0.0001 | <0.0001 |
| $I_2$ | 2 | <0.0001 | <0.0001 | <0.0001 | <0.0001 | <0.0001 | <0.0001 |
|  | 6 | <0.0001 | <0.0001 | <0.0001 | <0.0001 | <0.0001 | <0.0001 |
|  | 10 | <0.0001 | 0.0001 | <0.0001 | <0.0001 | <0.0001 | <0.0001 |
|  | 14 | <0.0001 | <0.0001 | <0.0001 | <0.0001 | <0.0001 | <0.0001 |
| $I_3$ | 2 | <0.0001 | 0.003 | <0.0001 | <0.0001 | 0.9249 | <0.0001 |
|  | 6 | <0.0001 | 0.0114 | <0.0001 | <0.0001 | <0.0001 | <0.0001 |
|  | 10 | 0.0009 | <0.0001 | 0.0268 | <0.0001 | 0.206 | <0.0001 |
|  | 14 | 0.0082 | <0.0001 | 0.0003 | <0.0001 | 0.1825 | <0.0001 |

**Table 6.** Statistics of C_RMS_ between kernel diameters. Two-sided permutation test with 10,000 repeats.

| Group |  | 2° vs. 6° | 2° vs. 10° | 2° vs. 14° | 6° vs. 10° | 6° vs. 14° | 10° vs. 14° |
| --- | --- | --- | --- | --- | --- | --- | --- |
| $I_1$ | Upper UV | <0.0001 | <0.0001 | <0.0001 | <0.0001 | <0.0001 | <0.0001 |
|  | Upper G | <0.0001 | <0.0001 | <0.0001 | <0.0001 | <0.0001 | <0.0001 |
|  | Lower UV | <0.0001 | <0.0001 | <0.0001 | <0.0001 | <0.0001 | <0.0001 |
|  | Lower G | <0.0001 | <0.0001 | <0.0001 | <0.0001 | <0.0001 | <0.0001 |
| $I_2$ | Upper UV | <0.0001 | <0.0001 | <0.0001 | <0.0001 | <0.0001 | <0.0001 |
|  | Upper G | <0.0001 | <0.0001 | <0.0001 | <0.0001 | <0.0001 | <0.0001 |
|  | Lower UV | <0.0001 | <0.0001 | <0.0001 | <0.0001 | <0.0001 | <0.0001 |
|  | Lower G | <0.0001 | <0.0001 | <0.0001 | <0.0001 | <0.0001 | <0.0001 |
| $I_3$ | Upper UV | <0.0001 | <0.0001 | <0.0001 | <0.0001 | <0.0001 | <0.0001 |
|  | Upper G | <0.0001 | <0.0001 | <0.0001 | <0.0001 | <0.0001 | <0.0001 |
|  | Lower UV | <0.0001 | <0.0001 | <0.0001 | <0.0001 | <0.0001 | <0.0001 |
|  | Lower G | <0.0001 | <0.0001 | <0.0001 | <0.0001 | <0.0001 | <0.0001 |

The differences in JSD between the upper and lower visual field likely reflect differences in chromatic contrast: When plotting green *C_RMS_* as a function of UV *C_RMS_*, the data points for the lower visual field tended to be distributed tightly along the identity line (e.g. Fig. 3d_1_,e_1_, bottom), indicative of a high correlation between the channels and, thus, low chromatic contrast. For the upper visual field, however, the *C_RMS_* distributions for UV compared to green were broader and shifted towards higher values, in particular for larger RF kernel diameters (e.g. Fig. 3e_1_, top), suggesting a lower channel correlation and high chromatic contrast, respectively.

In summary, we found that except for bright scenes, UV-green chromatic contrast was higher in the upper compared to the lower visual field, particularly for the large RF kernel diameters (Fig. 3g). Taken together, this suggests that the natural environment of mice above the horizon is rich in chromatic information, which may preferentially drive (colour-opponent) RGCs with large RFs.

### Natural scenes are biased towards dark contrasts

It has been reported that the contrast distribution of (monochromatic) natural scenes is biased towards dark (negative) contrasts and that this bias is mirrored in the higher proportion of Off vs. On responding neurons in the early visual system (Ratliff et al., 2010; Wang et al., 2015; Xing et al., 2010). Therefore, we went on to study the distribution of dark and bright contrasts in the recorded scenes (Fig. 4). To measure the contrast polarity distribution (“On-Off contrast”, *C_On−Off_*) in each channel, we convolved the crops separately with the center and surround of difference of Gaussian (DOG) kernels of various diameters (*d_RF_*, 2° to 14°) and computed the Michelson contrast between center and surround (Ratliff et al., 2010) (Fig. 4a-c). Like for *C_RMS_*, we analysed the distributions of *C_On−Off_* separately in both chromatic channels for the mean intensity groups (Fig. 4d-f).

**Figure 4.**
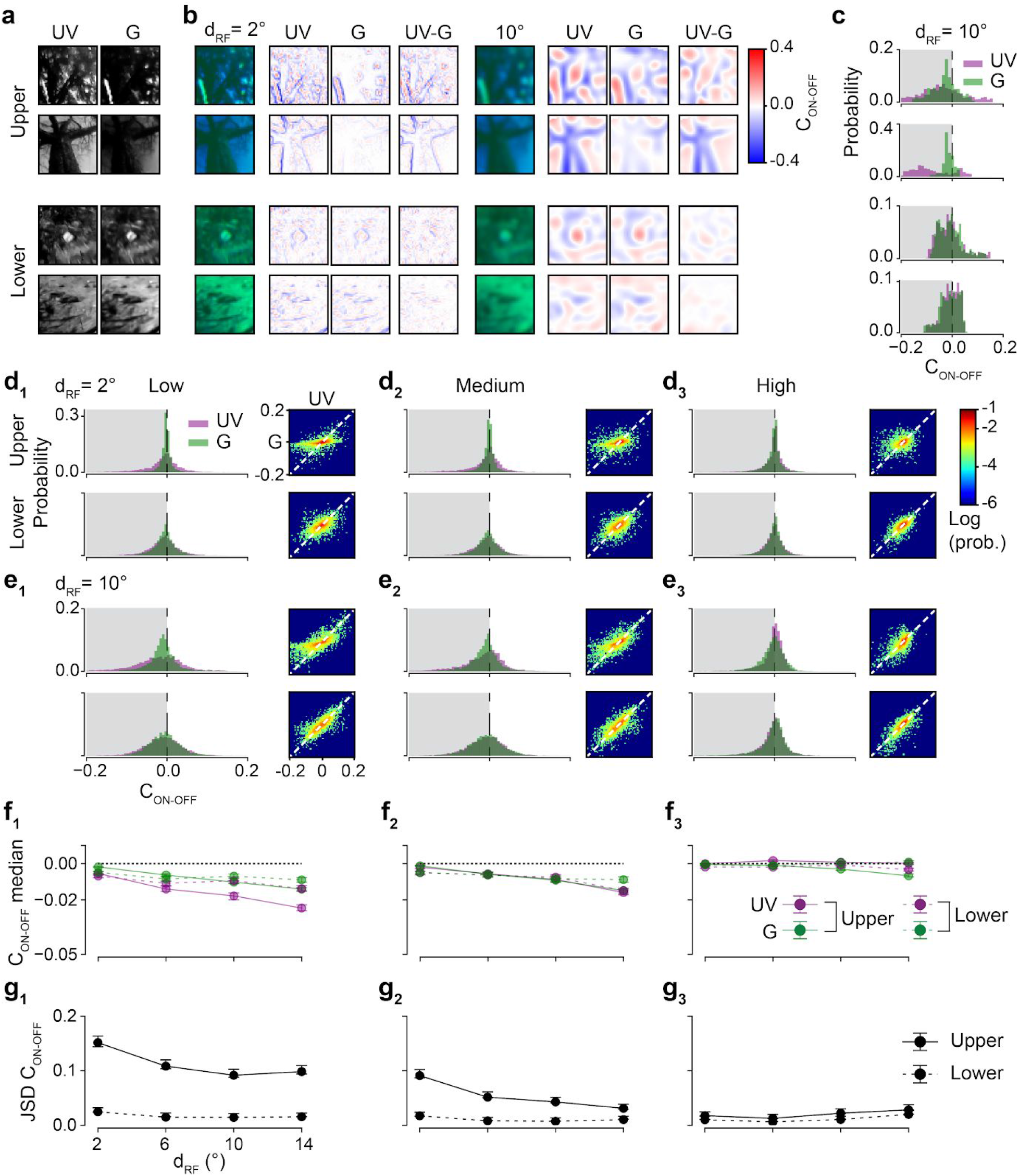
The upper visual field is biased towards negative contrast. **a**, Example image crops from upper and lower visual field (same as in Fig. 3a). **b**, images from (a) after application of On-Off filter kernel (Methods) with different RF diameters, d_RF_, with each of the two columns showing the UV-green overlay (left), UV (centre) and green (right) channel maps visualizing On-Off contrast (C_On−Off_). **c**, Distributions C_On−Off_ for UV and green for a medium-sized d_RF_ of 10°. **d,e**, 1D (left) and 2D histograms for C_On−Off_ of all n=1,500 image crops from randomly picked frames out of n=15 movies) for the three intensity groups (from Fig. 2d,e) and d_RF_=2° (d), d_RF_=10° (e). **f**, Median C_On−Off_ (± 0.1 SD) as a function of d_RF_ for UV and green channels in the upper (solid lines) and lower (dashed) visual field; for the three intensity groups. **g**, Like (f) but with Jensen-Shannon divergence (JSD) between C_On−Off_ distributions (cf 2D plots in (d,e)) of the two chromatic channels as a function of d_RF_. Error bars represent 2.5 and 97.5 percentiles with bootstrapping (see Methods). Note that for larger RF kernel sizes, the distribution mode was slightly shifted to negative values (e.g. (e)), yet, when pooling images across chromatic channels, visual fields and intensity groups, the distribution mode of all four kernel sizes was at zero (data not shown), in line with earlier observations (Ratliff et al., 2010).

In crops with low and medium mean intensities, we found the *C_On−Off_* distributions — despite being wide — to be skewed to negative values, particularly in the upper visual field and for larger RF kernel diameters (for examples, see Fig. 4a-c; for distributions, median and JSD, see Fig. 4d-g; for statistical comparisons, see Tables 7, 8). The dark bias in the upper visual field scenes resonates with results from an earlier study (Baden et al., 2013), which showed that mouse cone photoreceptors that survey the sky preferentially encode dark contrasts, suggesting a neural representation bias that starts already at the cone level.

**Table 7.**
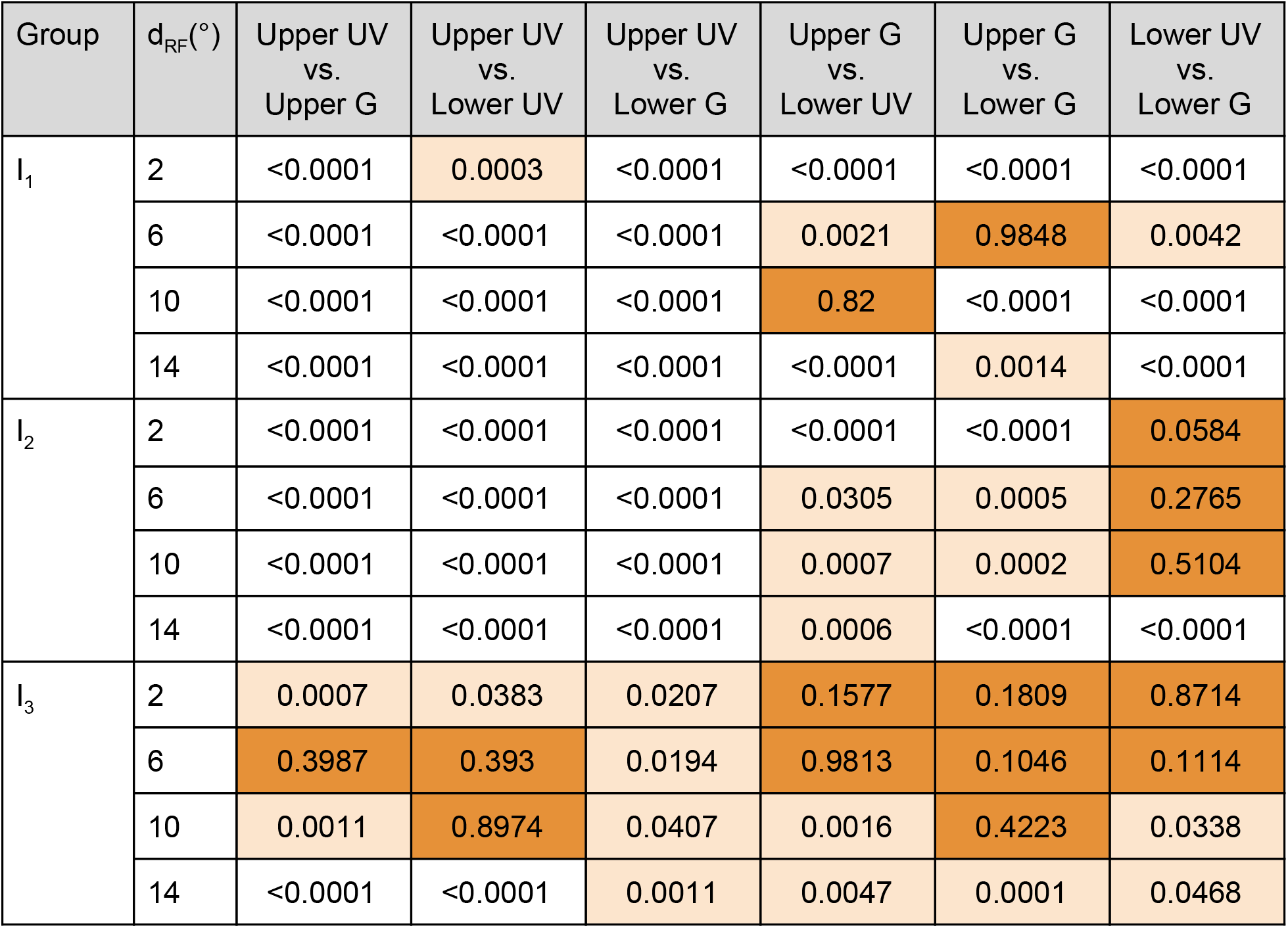
Statistics of C_On−Off_ between chromatic channels and upper vs. lower visual field as a function of kernel diameter. Two-sided permutation test with 10,000 repeats. White, light orange and dark orange indicating p < 0.0001, 0.0001 ≤ p < 0.05 and p ≥ 0.05.

**Table 8.**
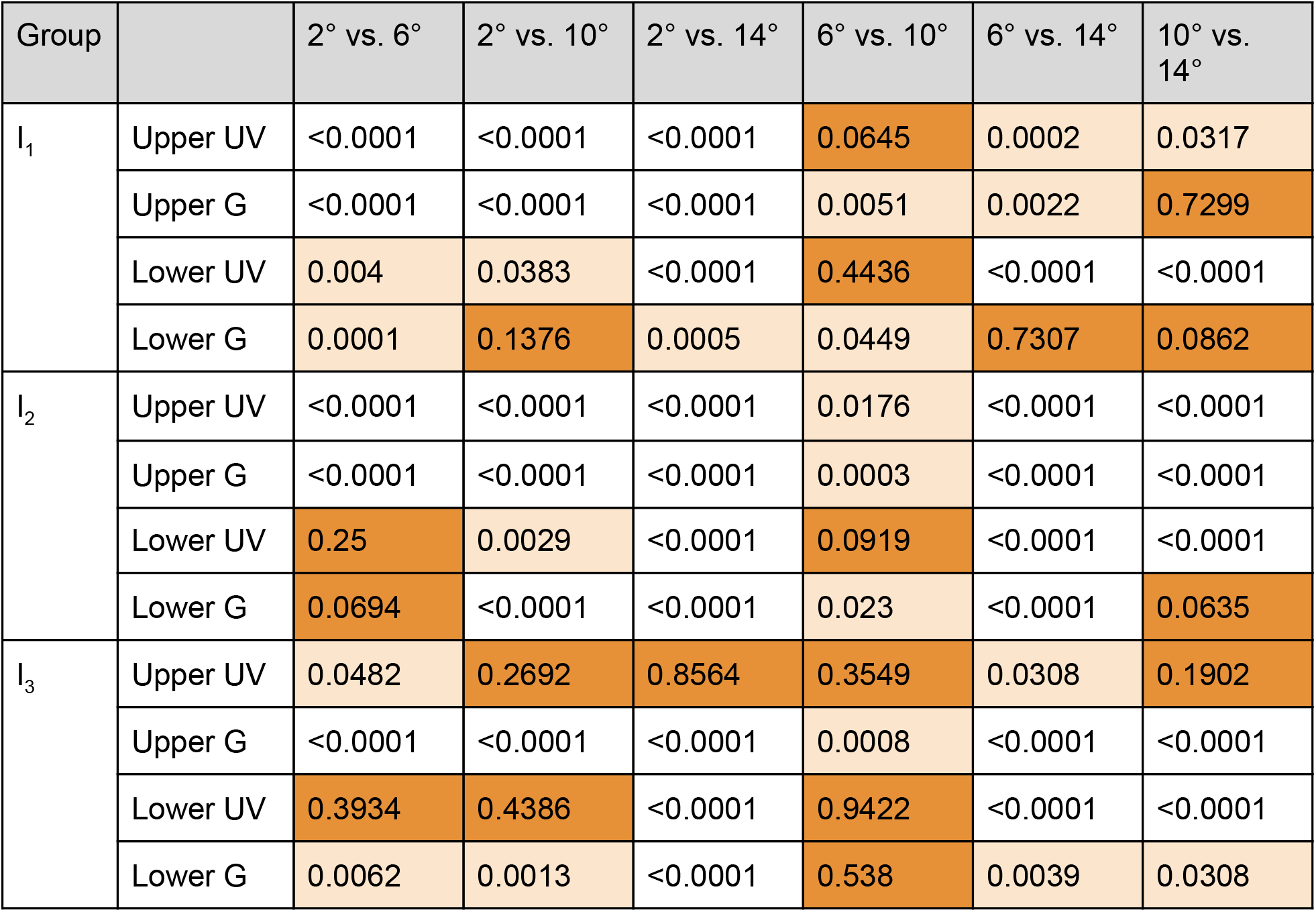
Statistics of C_On−Off_ between chromatic channels and upper vs. lower visual field as a function of kernel diameter. Two-sided permutation test with 10,000 repeats. White, light orange and dark orange indicating *p* < 0.0001, 0.0001 ≤ *p* < 0.05 and p ≥ 0.05

Hence, we next asked if such a neural representation bias is also present at the retinal output level, where we would expect a bias towards Off responses in ventral, large-RF RGCs. To test this prediction, we used a published dataset of RGC responses recorded in the ventral retina (Baden et al., 2016). We extracted the On-Off index (On, *OOi* > 0; Off, *OOi* < 0), which indicates if a cell responds more strongly to light-on or light-off transitions, and the RF diameter (*d_RF_*) for 2,380 RGCs (Suppl. Fig. S3a,b; see also Methods). We found that, indeed, Off cells with large RF (*d_RF_* > 8°, equivalent to >240 μ*m* in diameter) displayed a dark-biased *OOi* (Suppl. Fig. S3c) and were more frequent than their On cell counterparts (Suppl. Fig. S3d). This finding is consistent with the skewed distributions towards dark contrasts in our camera footage, suggesting retinal circuit adaptations that enable exploiting the dark bias present in natural scenes above the horizon.

We also compared the *C_On−Off_* distributions in the two channels and found chromatic On-Off contrast (quantified as *JSD*; Fig. 4g) to be systematically higher in the upper visual field, with the exception of the high mean intensity group. Moreover, like for the analysis of *C_RMS_*, the *C_On−Off_* distribution seemed broader for UV, supporting the idea that the mouse’s UV channel provides the animal with more nuanced information about the sky region than the green channel.

### UV channel for predator detection?

The high contrast available in the upper visual field and, specifically, in the animal’s UV channel may support detection of aerial predators (discussed in Cronin and Bok, 2016). However, so far, we focused on footage recorded during daytime and while mice are active during the day (Hut et al., 2011), they also forage from dusk till dawn. We thus asked if UV sensitivity was also useful for detecting objects in the sky at twilight. Interestingly, previous measurements of the spectral composition of sunlight over the course of a day revealed an overrepresentation of short wavelengths in an approx. one-hour period during twilight (Hut et al., 2000; Johnsen et al., 2006). This effect is due to absorption by ozone, together with the scattering of sunlight in the upper atmosphere, resulting in a blueish twilight sky (discussed in Johnsen et al., 2006). Since the ratio between 360 and 520 nm – near the sensitivity peaks of mouse opsins – was reported to increase at twilight (Hut et al., 2000), we recorded with our mouse-camera before sunrise and after sunset (Fig. 5). Because of the camera chip’s relatively low light sensitivity, we recorded movies in the direction of the sun from a fixed camera position (Fig. 5a).

**Figure 5.**
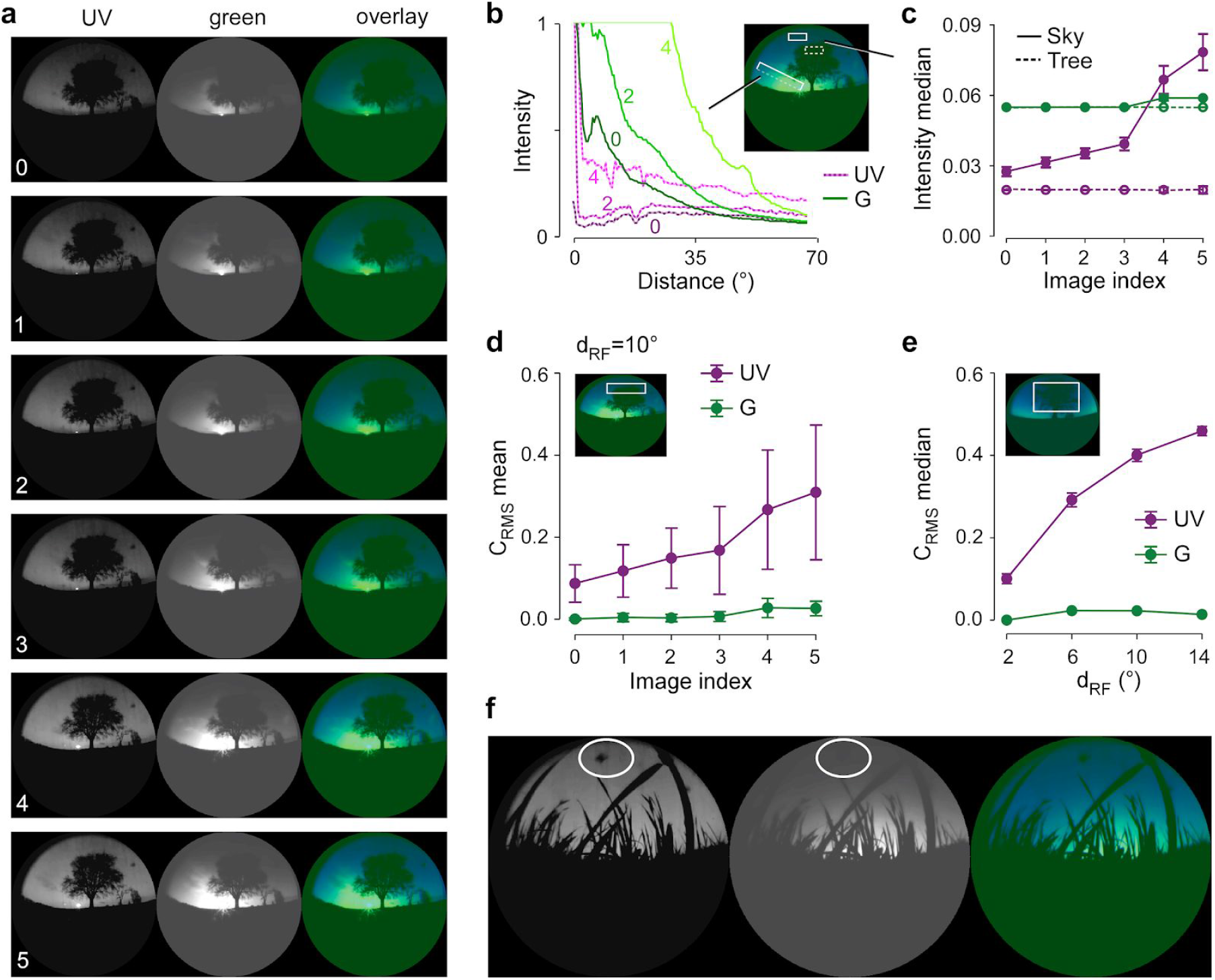
UV channel might facilitate detection of dark shapes in the upper visual field also at dusk and dawn. **a**, Representative scenes recorded with the mouse-camera around sunrise. **b**, Intensity profiles along a line (dashed, see inset) starting at the sun for images from (a). **c**, Median intensities in two image crops (see inset in (a); dashed box, tree; solid box, sky) as function of image series index. **d**, C_RMS_ (mean) in image crop at the edge of a tree (rectangle in inset) as function of image index. **e**, C_RMS_ (median) in image crop placed on the tree (rectangle in inset) as function of receptive field (RF) kernel diameter (d_RF_). **f**, Image showing an approaching black drone mimicking a bird of prey. Panels recorded at dawn (a-d) and dusk (e,f). Error bars in (c,d) and (e) representing 2.5 and 97.5 percentiles with bootstrapping.

First, we explored for the example frames the intensity profiles along a 70° arc starting at the sun’s position (Fig. 5b). In the vicinity of the sun, the intensity in the green channel was always higher than that in the UV channel, whereas further away from the sun, UV and green reached similar intensity levels (Fig. 5b; e.g. profiles from image #4 for >55°), in line with the observations by Hut et al. (2000). Moreover, for distances larger than approx. 5° from the sun, the UV intensity profile was much flatter than that of green, resulting in a more homogeneous UV illumination of the sky. Finally, in the sky’s dome, the intensity rose faster over the course of the sunrise in the UV compared to the green channel (Fig. 5c). Akin to daytime scenes, *C_RMS_* was significantly higher in the UV than the green (Fig. 5d; *p* < 0.0001, for all images, two-sided permutation test, *n* = 10,000 repeats) and increased with kernel diameter (Fig. 5e).

Together, these characteristics render the UV channel suitable for object detection, in particular for dark objects on top of a relatively bright (twilight) sky. We tested this hypothesis with a black drone mimicking an approaching aerial predator (Fig. 5f). As predicted, the drone was much more easily detectable in the UV vs. green channel, suggesting that this also holds for birds preying on mice during twilight, and therefore may contribute to increasing the species’ survival chances in their natural habitat (Suppl. Video V3).

### Autoencoder model predicts colour opponency in the ventral retina

So far, we characterized the contrast statistics of scenes from the mouse natural environment and found significant differences in chromatic contrast between upper and lower visual field. Next, we wanted to explore if these differences can give rise to colour opponency and, hence, shape neural representations. Recently, Ocko and colleagues (2018) trained a convolutional autoencoder (CAE; Ballard, 1987; Hinton and Salakhutdinov, 2006) model to reconstruct pink (1/*f*) noise and showed that the model learned spatial filters with center-surround organisation resembling the RFs of different RGC types. Thus, we next asked if such CAE models can also learn colour opponent RFs (Fig. 6). Autoencoder models feature a “bottleneck” that forces them to learn a “compressed” representation of the input images. This conceptually resembles the early visual system, where the information flow from the retina to downstream targets in the brain is constrained by bandwidth limitations (Gjorgjieva et al., 2014; Perge et al., 2009; Van Essen et al., 1991). We optimized our CAE model to yield good reconstructions under the qualitative constraint that the learned filters resemble smooth, centre-surround RFs (Suppl. Fig. S4). We then tested if colour-opponent kernels emerge in a CAE trained with our UV-green scene crops, and whether the total number of such kernels differed between the upper vs. lower visual field.

**Figure 6.**
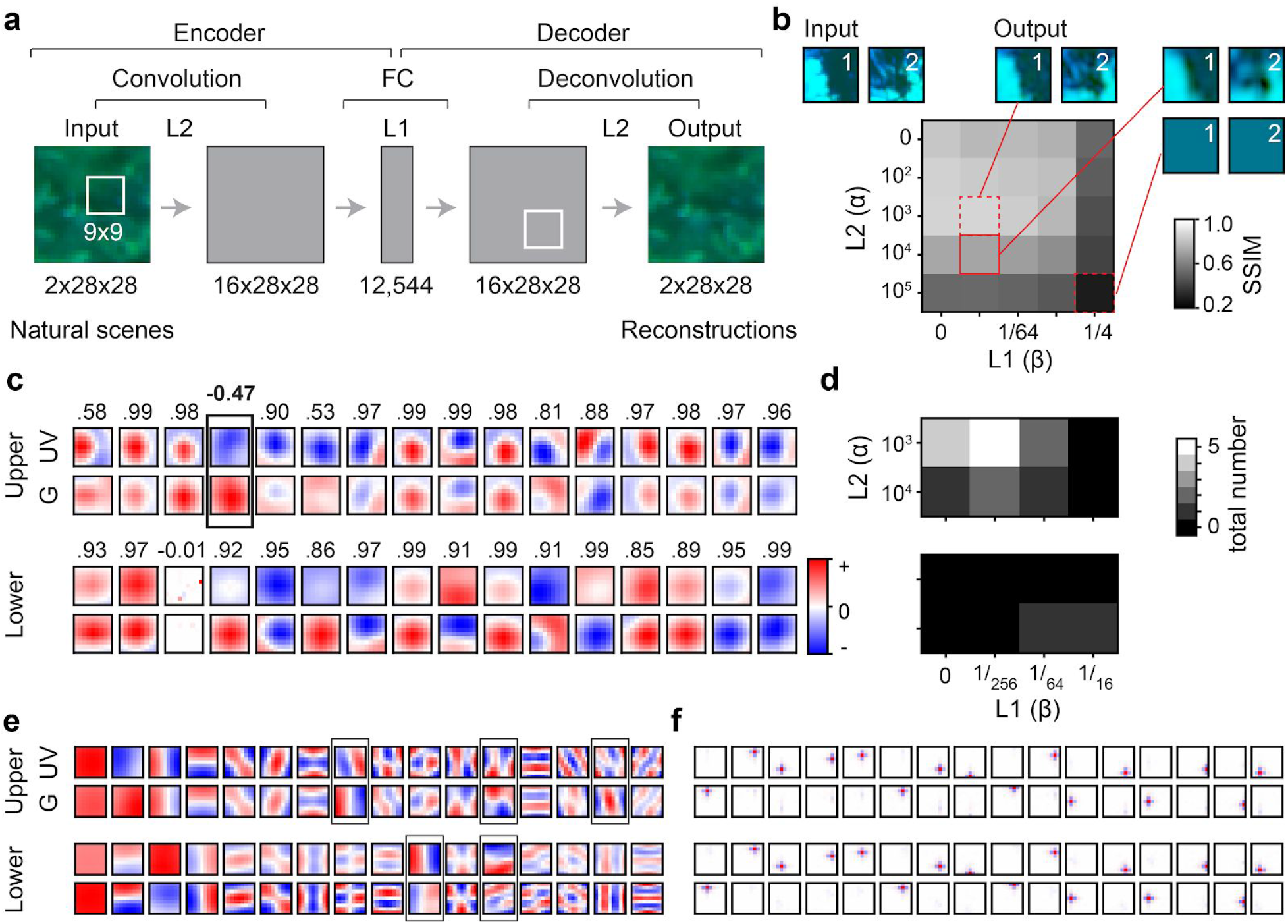
Convolutional autoencoder (CAE) model learns colour opponent kernels from upper visual field footage. **a**, Architecture of the CAE mode (FC, fully-connected; numbers indicate feature shapes; for details, see Methods)l. **b**, Reconstruction performance measured as structural similarity (SSiM) under different regularization strengths α (L2) and β (L1), with two example images (input; top-left) and the corresponding CAE outputs depicted for certain combinations of α and β (red boxes). **c**, Convolutional kernels learned by the model from upper (top) and lower (bottom) visual field scenes (for α=10^4^, β=1/256; see solid red box in (b); correlation indicated above each kernel pair; p<0.05 for all kernel pairs except third one in bottom row (p=0.9)). Example for colour-opponent kernel indicated by box. **d,** 2D histogram showing total number of colour-opponent kernels per random seed learned from upper (top) and lower (bottom) visual field scenes (n=10 random seeds). **e,f**, 16 kernels for UV and green, each for upper and lower visual field crops; same presentation and dataset as in (c). In (e), first kernels from PCA (Methods), with colour-opponent kernels indicated by boxes. The colour opponency index (COi; cf. Suppl. Fig. S2 and Methods) of the upper and lower visual field is 0.0393 and 0.0182, respectively. In (f), randomly selected kernels from ZCA whitening (Methods). Here, all 162 kernels (9×9 ×2 channels; cf. (a)) were colour-opponent for the upper visual field, while only 143 were colour opponent for the lower visual field.

Classically, autoencoders are encouraged to learn an efficient encoding of the input by reducing the number of neurons in the hidden layer. Instead, we here encouraged redundancy reduction of the input by adding Gaussian noise and imposing an L1 penalty (β) to the activations of the encoder’s output (Doi and Lewicki, 2007; Field, 1994; Ocko et al., 2018; for details, see Methods; van Rossum et al., 2003). To encourage smooth kernels akin to those measured in early visual neurons (e.g. Hubel and Wiesel, 1959; Marr and Hildreth, 1980), we additionally used L2 regularization (*a*) on the convolutional and deconvolutional layers, effectively constraining the norm of the weights (Vincent and Baddeley, 2003).

We trained the CAE model to minimize the mean squared error (MSE) for reconstructing images from the upper and lower visual field, separately for different regularizations and different random initialization seeds. We then quantified the model’s performance in reconstructing the input images (Fig. 6b) using structural similarity as metrics (SSIM; Wang et al., 2004). Qualitatively, we searched for combinations of α (L2) and β (L1) that used smooth centre-surround-like kernels in the presence of a bottleneck to reliably yield good reconstruction performance (*SSIM* ≥ 0.6; Fig. 6c). For these parameter combinations, we ran the model with *n* =10 random seeds and determined the number of colour-opponent spatial kernels (Fig. 6d). Kernels were considered colour-opponent if their UV and green channel were negatively correlated (*p* < 0.05; Pearson correlation; cf. Fig. 6c).

We found that colour-opponent kernels were significantly more frequent in CAEs trained with scenes from the upper vs. the lower visual field (*p* = 0.012, permutation test; Fig. 6d). This predicts that in systems with an information bottleneck, UV-green colour-opponent kernels preferentially emerge when encoding visual scenes above the horizon.

We compared our CAE model with classical approaches that explain the unsupervised emergence of colour-opponency (Buchsbaum and Gottschalk, 1983) or centre-surround RF structure from natural stimuli (Bell and Sejnowski, 1997; Graham et al., 2006). We first applied PCA (Suppl. Fig. S2) to 9×9 image patches randomly drawn from the same set of image crops, and found that crops from the upper visual field have a higher *CO_i_* (higher variance along the colour-opponent dimension; cf. Suppl. Fig. S2) than ones from the lower visual field (Fig. 6e). Unlike the CAE, the PCA kernels did not resemble the centre-surround RFs known in RGCs. Similarly, we applied zero-phase component analysis whitening (ZCA; Methods; Bell and Sejnowski, 1997). Also here, we found more colour-opponent kernels with images from the upper visual field (Fig. 6f). However, while these kernels were centre-surround, they were small and mainly different in spatial position.

In summary, all three unsupervised models confirm that specific chromatic statistics in the upper visual field may be sufficient to drive the emergence of colour-opponent spatial RFs.

## Discussion

Using a custom-build camera with spectral channels matching the UV and green sensitivities of mouse photoreceptors, we captured scenes from the natural visual environment of mice. Statistical analyses of these scenes revealed that chromatic contrast was stronger above the horizon, and that such contrast would preferentially drive large RFs. This enrichment of chromatic contrast in the upper visual field might underlie the preferential emergence of colour-opponent RFs found in the ventral retina, as suggested by a convolutional autoencoder model trained to represent our UV-green scenes. Furthermore, we found the upper visual field, and in particular the UV channel, to be biased towards dark contrast, which supports the view that UV could be an important source of information about dark objects (i.e. birds) against a brighter sky. Together, our findings lend further support to the idea that retinal circuits have evolved to optimally process natural scene statistics (reviewed in Baden et al., 2020).

### Recording natural scenes

Because we aimed at a camera that was portable, low-cost, and easily reproducible, we opted for the well-documented OV5647 camera chip of the second-generation Raspberry Pi camera. The main disadvantage of this choice was the relatively low light sensitivity of the chip, which restricted the recordings of dynamic natural scenes to daytime and twilight. With other camera chips — e.g. the new Raspberry Pi High Quality camera or smartphone cameras optimized for low-light recordings — a more nighttime-enabled mouse-camera, potentially also with a larger dynamic range, may be in reach. A further improvement may be opsin-matched filters (Tedore and Nilsson, 2019). Another interesting direction would be to miniaturize the camera to allow it to be mounted on a mouse while it roams its environment. Current head-mounted cameras for mice are restricted to grayscale or RGB (Froudarakis et al., 2014; Meyer et al., 2018, 2020; Michaiel et al., 2020) and therefore miss the UV channel altogether (see discussion in Franke et al., 2019). For an indoor environment lacking UV illumination this may not be relevant, but, as demonstrated in our study, when the settings approach the natural habitat of mice, a UV channel is a rich source of potentially behaviour-relevant visual information.

### Mouse camera movies as natural stimuli

Stimuli used in vision research cover a broad spectrum, ranging from artificial stimuli, like gratings and noise (reviewed in Rust and Movshon, 2005) or screen-rendered 3D objects (Froudarakis et al., 2020; Zoccolan et al., 2009), to more naturalistic ones (e.g. Baddeley et al., 1997; Betsch et al., 2004; Froudarakis et al., 2014). We argue that the movies recorded here represent suitable natural stimuli to probe mouse vision for the following reasons: (*i*) They were recorded outdoors in fields where mice can be seen during the day, and (*ii*) contain the spectral bands perceivable by mice. If presented on a UV-capable stimulator, they should drive the mouse visual system more efficiently than gray-scale stimuli on a standard monitor (discussed in Franke et al., 2019). (*iii*) Because our camera was mounted on a gimbal, the movies approximate well the input that reaches the mouse’s eye, since a large fraction of eye movements in mice serve to stabilize the retinal image in the presence of head movements (Meyer et al., 2020). A first validation of these stimuli could be to test if they allow separating anatomically defined RCG types that so far could not be reliably disambiguated based on their responses to standard synthetic stimuli (Baden et al., 2016).

### Natural scene representations in the mouse early visual system

#### Retinal opsin gradient, colour opponency and colour vision

Like many vertebrates, mice display regional opsin co-expression in cone photoreceptors (reviewed in Peichl, 2005): Specifically, ventral M-cones co-express S-opsin (Applebury et al., 2000; Baden et al., 2013; Röhlich et al., 1994; Szél et al., 1992), a challenge for a colour vision mechanism that relies on comparing signals from spectrally distinct cone types. Yet, our scene analysis showed chromatic contrast to be higher in the upper visual field, which is “viewed” by the heavily S-opsin dominated ventral retina. Moreover, the results from our autoencoder model suggests that the chromatic information in the upper visual field may have driven the presence of colour-opponent cells in the ventral retina over the course of evolution. In line with this prediction, a recent neurophysiological study showed that colour-opponent RGCs are indeed more frequent in the ventral retina (Szatko et al., 2020). At least some of these colour-opponent RGCs rely on antagonistic centre-surround RF mechanisms, with the RF centre dominated by signals from UV-biased S/(M)-coexpressing cones and the green-biased RF surround being mediated by rods (Joesch and Meister, 2016; Szatko et al., 2020) and/or by long-ranging inputs from M-cones in the dorsal retina (Chang et al., 2013). We also found more colour-opponent kernels for the upper vs. the lower visual field when employing a similar definition of centre-surround opponency as Szatko and coworkers (Szatko et al., 2020). Interestingly, these mechanisms should profit from pooling signals from a large surround, which resonates well with our finding that larger RF cells should be better suited to pick up chromatic contrast. Further support for a crucial role of the ventral retina in promoting colour vision comes from behaviour: By probing the visual field of mice with chromatic stimuli, Denman et al. (2018) found that the animals succeeded in a colour change detection task for stimuli presented in the upper visual field.

If this chromatic regionalization of the retina was indeed evolutionary advantageous, why is it not more widespread among critters in similar habitats? For instance, some members of the subfamily Murinae, like the steppe mouse (*Mus spicilegus*), shares S/M opsin regionalization with the house mouse (*Mus musculus*), whereas others lack it (*Apodemus sylvaticus*) or lost the S-opsin completely (*Apodemus microps*) (Szel et al., 1994). Interestingly, these Apodemus species live in shrubberies, where a “bird-in-the-sky” detector may not be as useful as in relatively open spaces, as inhabited by *M. spicilegus*. Hence, spectral retinal regionalization in some mice may reflect an environmental adaptation — akin to the UV-sensitive “strike zone” in the ventral retina of zebrafish larvae, which helps the animal detect UV-reflecting prey and avoid potential predators (large dark shapes) in the upper visual field (Zimmermann et al., 2018). Curiously reminiscent of this “strike zone”, Nadal-Nicolás and colleagues (2020) recently found a S-cone “hotspot” in the ventral periphery of the mouse retina.

#### UV vision

UV sensitivity is thought to play an important role in the behaviour of many vertebrates and invertebrates, including navigation, orientation, communication, foraging, as well as predator and prey detection (reviewed in Cronin and Bok, 2016). In rodents, UV vision has been mainly discussed in the context of communication via urine marks (Chávez et al., 2003; Joesch and Meister, 2016) and predator detection (e.g. Baden et al., 2013), but it may also play a role in foraging with respect to enhanced chromatic contrast of foliage (Tedore and Nilsson, 2019) and fruit (Altshuler, 2001). Our results shed light onto why mice specifically profit from UV sensitivity for predator detection: Consistent with previous studies (Baden et al., 2013; reviewed in Cronin and Bok, 2016), our results suggest that during daylight, when S-cones are at their prime, UV sensitivity enhances the detection of dark silhouettes against the bright sky. At twilight, the spectrum becomes more dominated by short wavelengths (e.g. increase in ratio between 360 and 440 nm, see Hut et al., 2000). From our footage, we estimated the intensity of the sky’s dome around sunset in the UV channel to be equivalent to ~ 300 *R*s*^−1^ per cone. This is well above the S-cone’s threshold of ~20 *R*s*^−1^ per cone (Naarendorp et al., 2010), even when including some UV-filtering by the mouse optics (33% transmission at 360 nm; Henriksson et al., 2010), and above the levels of ~ 100 *R*s*^−1^ per rod below which rods are thought to take over (Rodieck, 1998). Moreover, recent V1 recording and modeling results suggest that around sunrise, mouse vision becomes quickly dominated by cones (Rhim et al., 2020). Under these conditions, S-cones, which are also less noisy than M-cones (Ala-Laurila et al., 2004), may indeed contribute to detecting dark shapes in front of the sky. Finally, at even lower intensities, when the rod photoreceptors dominate (Naarendorp et al., 2010), UV light from the sky may still play a role, because rhodopsin — like any opsin — features a secondary sensitivity peak (“beta band”) around ~350 nm (Govardovskii et al., 2000). Therefore, it is conceivable that at low light levels, mice may perceive near-UV via the rods (discussed in Cronin and Bok, 2016).

#### Bias towards dark contrasts

Our analysis of mouse natural scenes is consistent with previous studies, showing that the distribution of contrast in natural scenes is biased towards dark contrasts (Cooper and Norcia, 2015; Ratliff et al., 2010). The dark bias in our natural scene footage was more prominent in the upper visual field and specifically in the UV channel, which resonated well with earlier retinal recordings showing that light-off steps are particularly faithfully encoded by ventral cones (Baden et al., 2013). Similar to previous findings that describe an overrepresentation of Off information in the visual system (Kremkow et al., 2016; Schröder et al., 2020; Yeh et al., 2009), we also found that dark contrasts elicited stronger responses than bright contrasts, particularly in ventral RGCs. Interestingly, the dark bias was most prominent in large-RF RGCs, which ties into an ongoing controversy if Off cells are expected to feature large (e.g. Cooper and Norcia, 2015) or small RFs (see discussion in Mazade et al., 2019; e.g. Ratliff et al., 2010). We currently lack the corresponding RF data for dorsal RGCs, and therefore the question remains if the balanced contrast distribution in the lower visual field is also reflected in a more balanced distribution of On/Off preferences.

#### Eye movements in the context of a horizontally regionalized visual field

As discussed above, several lines of evidence suggest that the dorsal and ventral mouse retina are functionally specialized. Such dorso-ventral regionalization is expected to be advantageous only if it is ensured that the upper and lower visual field is reliably projected onto the ventral and dorsal retina, respectively. This requires a different visual orienting strategy compared to primates, who scan the visual environment with their foveas using head-motion independent eye movements (Baden et al., 2020; reviewed in Land, 2015). Eye movements in mice are tightly coupled to head movements, and often serve to stabilize the retinal image with respect to the ground (Meyer et al., 2018, 2020; Michaiel et al., 2020). This presumably ensures that the temporo-nasal retinal axis is aligned with the horizon and would thus allow the mouse retina with its dorso-ventral specialization to make use of the respective differences in natural scene statistics.

### Are natural scene statistics encoded in retinal circuits?

Autoencoders have long been used to learn efficient representations by restricting the coding capacity of the hidden layer(s) (Hinton and Salakhutdinov, 2006; Kramer, 1991). Therefore, they are a natural choice for exploring feature transformations in the early visual system, such as between retina and primary visual cortex (V1), where visual information has to pass the optic nerve, posing a severe bottleneck. Recently, Ocko and colleagues (2018) trained a convolutional autoencoder with pink (1/*f*) noise, which mimics the distribution of spatial frequencies in natural scenes (Field, 1987). They found that it learned spatial filters that resembled centre-surround RFs of a subset of primate RGC types. We used a similar CAE architecture trained with natural UV-green images, which learned, in line with Ocko et al. (2018), not only spatial filters resembling diverse RGC RFs, but also colour-opponency (Buchsbaum and Gottschalk, 1983). While other approaches, such as PCA and ZCA whitening, also revealed colour-opponency and, in case of ZCA whitening, centre-surround structure, the CAE was unique in the diversity of resulting spatial filters reminiscent of different RGC types. In the future, it would be informative to investigate which types of regularization mimic the specific constraints represented by bottlenecks in the early visual system (and why) and, by systematically manipulating the input images, what particular features give rise to colour opponency (discussed in Chalk et al., 2018). Also, it would be interesting to explore if the kernels learned by the CAE indeed help predicting RGC responses to natural or synthetic stimuli.

## Acknowledgments

We thank Tom Baden, Heiko Schütt, and Matthias Bethge for helpful discussions, and Gordon Eske for excellent technical assistance. This work was supported by the German Research Foundation (DFG) through Collaborative Research Centre CRC 1233 (project number 276693517); the European Union’s Horizon 2020 research and innovation programme under the Marie Skłodowska-Curie grant (agreement No 674901); and the Max Planck Society (M.FE.A.KYBE0004); the German Ministry of Research and Education (01GQ1002). The funders had no role in study design, data collection and analysis, decision to publish, or preparation of the manuscript.

## Author Contributions

Conceptualization Ideas: F.S., K.R., L.B., and T.E.; Methodology: Y.Q., Z.Z., L.B., and T.E., with input from D.K. and M.K.; Formal Analysis: Y.Q. with input from D.K., L.B., and T.E.; Investigation: Y.Q., Z.Z., K.S., with input from K.F., L.B. and T.E.; Writing – Original Draft: Y.Q., L.B., and T.E.; Writing – Review & Editing Preparation: Y.Q., Z.Z., D.K., K.S., M.K., K.F., L.B., and T.E.; Visualization: Y.Q., and T.E.; Supervision: L.B. and T.E.; Funding Acquisition: L.B., F.S., K.R., and T.E.

## Methods

### Camera design

As the front lens of our camera (Fig. 1a), we used a fisheye lens (*f* = 1.05 *mm*, *f*/2, LS-40180, Watterott, Germany) with a FOV of approx. 180°. Following an inverted periscope design, the lens was mounted at the bottom end of the camera, which allowed us to capture the scene from a distance of approx. 2-5 cm to the ground (Fig. 1b-f). After passing two relay lenses (L04, LB4879-A, *f* =35 *mm*; L05, LB4854-A, *f* =20 *mm*; both Thorlabs, German), which allowed us to transfer the fisheye lens’ FOV to the camera sensor, a silver mirror reflected the light towards a dichroic mirror (F48-442, AHF, Germany; reflection, > 90%, 350–442 *nm*; transmission, >90%, 460–650 *nm*; Fig. 1c) that reflected wavelengths shorter than approx. 440 *nm* towards the first camera sensor (CS1) and transmitted longer wavelengths towards the second camera sensor (CS2; Fig. 1a). An additional spectral bandpass filter in front of each camera chip (BP_G_, F47-510, >90%, 470-550 *nm*; BP_UV_, F37-424, > 90%, 350–419 *nm*; both AHF) restricted the light reaching the chips to the approx. spectral ranges relevant for mouse opsins (Fig. 1c). Because the spectral properties of dichroic filters (DM, BP_UV_, BP_G_) change with the incident angle of the light, we used relay lenses L1, L2 (LA4148-A, *f* =50 *mm*; Thorlabs, Germany) and L3 (LA4874-A, *f* = 150 *mm*; Thorlabs) to ensure that light passes the dichroic bandpass filters collimated before being focussed onto the camera chips. In addition, we added iris diaphragms (SM1D12C; adjusted to ~2 *mm* pinhole diameter) directly in front of the cameras to optimize depth-of-field and image contrast. For the parts list, including mechanical parts, see Table 1.

For each chromatic channel, we used a Zero spy camera module (EXP-R15-1309, EXP-TECH, Germany) connected to separate Raspberry Pi Zero W single-board computers (EXP-R12-143, EXP-TECH), which were powered by a 15 Ah USB power bank (Anker 1252011, Reichelt, Germany). Lenses and infrared filters had been removed from the camera modules. To increase the UV sensitivity of the chip in the UV pathways, we mechanically removed its RGB Bayer layer, following a procedure described by Wilkers et al. (2016).

#### Movie recordings

Movies were recorded onto the Raspberry Pi’s Flash memory card; movie capture was remote-controlled from a laptop connected to the Raspberry Pis either via USB cable or an ad-hoc Wifi network (VNC Viewer, RealVNC Ltd, UK). Camera parameters were fixed for all recordings (Table 2). To stabilize the camera during the recordings, we mounted it on a gimbal (Crane 2, Zhiyun, China; Fig. 1d,e). Since the camera weighed around 1 kg and its point of gravity was not centred, we added counterweights for the gimbal to work properly. When moving, we tried to maintain an azimuth angle of ~60° between the optical axis of the fisheye lens and movement direction, close to the azimuth angle of the mouse’ eye (Oommen and Stahl, 2008; Stabio et al., 2018; Sterratt et al., 2013).

#### Temporal alignment

Since we recorded with two camera chips simultaneously, the resulting movies needed to be temporally and spatially (next section) synchronized. For temporal alignment, we used LEDs mounted close to each camera chip (Fig. 1a) and flashed (200-ms pulses every 20 seconds) them as synchronisation markers. In addition, we manually checked the temporal alignment by comparing frames from the two channels (e.g. during fast movements).

#### Spatial alignment

Because we used optical rails to build the mouse-cam’s optical pathway, the UV and green channels captured almost the same scene. To account for potential spatial offsets and differences in image magnification, we used a homography matrix (*H*) to spatially relate a point (*x*_1_, *y*_1_)^*T*^ in the first channel to a corresponding point (*x*_2_, *y*_2_)^*T*^ in the other channel:

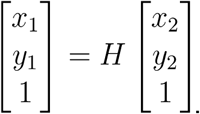

To this end, we first extracted at least 20 feature points from both channels using the scale-invariant feature transform (SIFT) approach (Lowe, 2004) and matched these feature points using the k-nearest neighbors algorithm. Next, *H* was determined by a random sample consensus algorithm (Fischler and Bolles, 1981), allowing us to project all pixels in the first channel (green) to those in the second channel (UV), with the same *H* typically working for all frames of a particular recording.

#### Spectral calibration

To map image pixel values (*v_Raw_*, 0…255) to absolute intensities (in [μ*W*]), we first intensity-calibrated LEDs peaking in the UV (λ_*Peak*_ = 360, 380, 400 *nm*; see Table 3; Suppl. Fig. S1a) or the green band (λ_*Peak*_ = 490, 525 *nm*) using a power meter (842-PE, Newport, Germany). Next we recorded images of these LEDs set to different intensities to determine the relationship between normalized pixel value (*v*, 0…1) and power (*P*) by fitting the data to *P* = *av^b^* + *c* (Suppl. Fig. S1b; for coefficients, see Table 2) using the Levenberg-Marquardt algorithm for least squares curve fitting. Ground-truth spectral images of the natural scenes were acquired with a scanning spectrometer, as described earlier (Baden et al., 2013). In brief, this custom-built device consisted of two servo motors that moved the fibre of a USB-spectrometer (STS-UV, Ocean Optics, Germany) to rasterize the scene with a resolution of ~10° of visual angle (Suppl. Fig. S1d,f, right).

To use the available range for both colour channels, our intensity correction linearly mapped an intensity range of 0.02 – 0.76 μ*W* (UV) and 0.37 – 6.56 μ*W* (green) to pixel values between 5 – 255 and 14–255, respectively (see Discussion). Note that for better visualisation, we applied gamma correction (*v*′ = *v*^1/*γ*^, with γ = 2.2; (Poynton, 2003)) to the images in the figures (e.g. Suppl. Fig. S1c,d vs S1f).

#### Statistical analysis of the natural scenes

A movie frame contained a circular FOV of ~180°, corresponding to 437 pixels along the diameter. To minimize the influence of potential chromatic and spatial aberrations introduced by the fish-eye lens, we focused on image cut-outs (“crops”; (53°)^2^, equivalent to (128 pixels)^2^ in size) from the central upper and lower visual field. For contrast analysis, we excluded image crops that contained more than 30% underexposed (*v_Raw_*(*G*) < 15, *v_Raw_*(*UV*) < 6) or overexposed (*v_Raw_*(*G*), *v_Raw_*(*UV*) > 254) pixels. We randomly sampled one image crop every 10 frames from all movies (Fig. 1b,e) until we had 1,500 crops for each upper and lower visual field. Next, the image crops were divided into three “intensity classes” (*I_low_, I_median_, I_high_*) by two percentiles (1/3 and 2/3) by their mean intensities.

Note that scene content varied somewhat with the image group, because scenes in or near the forest were usually dimmer (“low mean”) than those with open skies, bare ground and little vegetation (“high mean”). Since this bias was merely a side-effect of our mean intensity criterion, we refrained from linking statistics to scene content. In the future, it would be interesting to explore how scene content affects chromatic contrast statistics, for instance by classifying the images with pre-trained CNN models like VGG (Simonyan and Zisserman, 2014) before the statistical analyses. This could, for example, shed light on the question why contrast differences generally tended to decrease towards the high mean intensity group — that is, whether this is simply due to the limited dynamic range of the camera or in fact scene content-dependent. To determine *C_RMS_* and *C_On−Off_*, we randomly picked in each class for all images 10 locations per crop.

#### Root mean square (RMS) contrast

In psychophysical studies, *C_RMS_* is commonly used for estimating contrast in natural scenes (Mante et al., 2005) and defined as:

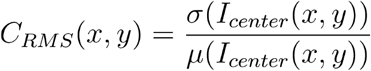

where σ(*I_Centre_*(*x, y*)) and μ(*I_Centre_*(*x, y*)) are standard deviation and mean, respectively, of the normalized pixel intensities contained in an image spot (“receptive field”; RF) centred at (*x, y*) within the image crop. Spot diameters (*d_RF_*) ranged from 2 to 14 degrees of visual angle.

#### On-Off contrast

We measured On-Off contrast (*C_On−Off_*) at a point (*x, y*) using a difference-of-Gaussians (DOG) kernel with a normalized denominator to restrict the value range to [−1, 1]:

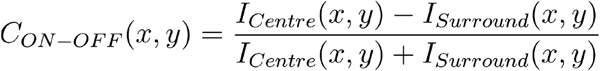

where *I_Centre_*(*x, y*) and *I_Surround_*(*x, y*) represent the summed pixel intensities after convolving the image with the centre and surround Gaussian kernels, respectively. The spatial relationship between centre and surround Gaussians were σ_*Surround*_ = 1.5σ_*Centre*_; the total DOG kernel size was 3σ_*Centre*_. Note that *d_RF_* was defined by the zero crossing radius of the kernel:

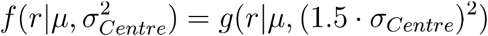

where 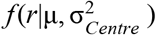 and 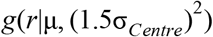 are centre and surround Gaussians, respectively. We ensure that the kernel response to a homogeneous input image 0 by setting *r* = 1.2σ_*Centre*_. A negative kernel response resulted in *C_On−Off_* < 0, indicating a negative contrast at this sample location. As for *C_RMS_, d_RF_* ranged from 2° to 14°.

#### On-Off index

To test if ventral mouse RGCs prefer dark contrasts, we reevaluated a published dataset with recordings of ventral RGCs (Baden et al., 2016). From this dataset, we extracted for all On (groups 1-9) and Off (groups 15-32) RGCs that passed the quality criterion (*Qi* >0.2; *n* = 2,380 cells) the On-Off index (cell_oo_idx, *OOi*; see below) and the RF diameter (rf_size, in [μm]). The latter was converted from [μm] into [°] of visual angle, assuming 1°≈30 μ*m* on the mouse retinal surface. The On-Off preference (*OOi*) of a cell and was defined as:

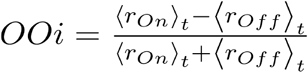

with *r_On_* and *r_Off_* defined as the activity during the response to the leading and the trailing edge of a bright-on-dark moving bar stimulus, respectively.

#### Comparing contrast distributions

To test if two contrast distributions originate from the same distribution, we performed a two-sided permutation test with 10,000 repeats to estimate the p-value. In addition, as a metric for similarity between the contrast distributions in the UV (*P_UV_*) and the green (*P_G_*) channel, we used Jensen-Shannon divergence (JSD):

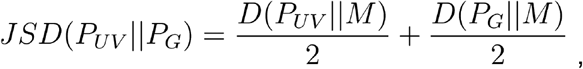

with the Kullback-Leibler (KL) divergence (*D*) defined as:

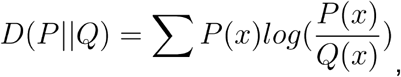

and *M* = 0.5(*P_UV_* + *P_G_*). Instead of KL divergence, we used JSD because it is symmetric and bounded (0…1 for log base 2).

### PCA and ZCA whitening

Buchsbaum and Gottschalk (1983) have shown that achromatic and chromatic visual channels can be obtained using principal component analysis (PCA). As an extension, zero-phase component analysis (ZCA) whitening was shown to decorrelate signals and learned centre-surround-like kernels (Bell and Sejnowski, 1997). Accordingly, we defined a covariance matrix *C* of the original centred data 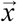. By applying transformation matrix *W*, we would get uncorrelated data 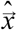

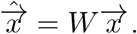

We performed PCA (without whitening) on the image crops using:

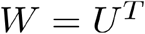

ZCA whitening was performed using

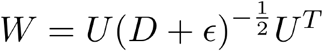

where *U* and *D* contain eigenvectors and eigenvalues of *C*, respectively, and ε = 10^−8^ for numerical stability.

We first applied PCA on intensities of chromatic channels. The eigenvector with two positive or two negative entries corresponds to an achromatic transformation and the other eigenvector with one positive and one negative entry corresponds to a chromatic (colour-opponent) transformation. We also applied PCA on 9×9 image patches. We defined the colour-opponent transformation when the kernels from the UV and green channels were negatively correlated (pearson correlation coefficient, *p* <0.05). In both cases, the colour opponency index (*CO_i_*), which represents the ratio between signal variance in the colour-opponent dimensions to the variance in all dimensions, was defined as:

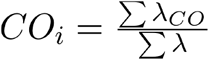

where λ_*CO*_ denotes the eigenvalues of the colour-opponent transformation, and λ all eigenvalues.

### Convolutional autoencoder model

We prepared datasets from the upper and the lower visual field separately. For both datasets, 10,000 image crops (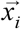 and *i* represents image index) with 56×56 pixels each were randomly picked and then rescaled to 28×28 pixels. These image crops met the same quality criteria (fewer than 30% of pixels under/over-exposure) as those for the statistical analysis. Among them, 9,000 image crops were used for training and the rest for validation and testing. The images were offset-corrected separately in each chromatic channel (by subtracting the channel’s mean intensity).

We implemented a simple convolutional autoencoder model (CAE; Fig. 6a) following (Ocko et al., 2018) using PyTorch (Paszke et al., 2019). The encoder contained a single convolutional layer (with weights denoted 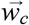) followed by a rectified linear unit (ReLU) function, one fully-connected (FC) layer and another ReLU function. Following Ocko et al. (2018), Gaussian noise with σ = 1 was added to the encoder output to restrict the channel capacity. The decoder contained one FC layer, one ReLU function, a single deconvolutional layer (with weights denoted 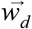), and a hyperbolic tangent (tanh) function to map back to the original data range (−1…1). We used 16 convolutional kernels with a filter size of 9×9 pixels for each chromatic channel, with zero-padded boundaries and without downsampling. Correspondingly, the deconvolutional kernels consisted of 9×9 pixel filters per input channel. Thus, the size of the activation tensor after the first convolution was 28×28×16 (height x width x channel), which was flattened into a 12,544 dimensional vector before it was fed into the FC layer. The two FC layers had the same input and output size. The loss function was defined as:

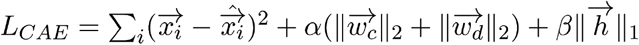

The first term is the MSE between prediction 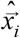 and ground truth 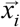, the second term is the L2 penalty (hyperparameter α) on the weights of the convolutional 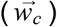 and deconvolutional 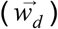 layers, and the third term is the L1 penalty (hyperparameter β) on the encoder output (see also Results).

The CAE models were trained for 100 epochs with 100 image crops in each mini-batch (learning rate, η = 10^−4^) using the Adam optimizer (Kingma and Ba, 2014) to minimize the loss functions. Image reconstruction performance of the CAE was estimated based on structural similarity (SSIM; Wang et al., 2004) and MSE (Fig. 6b; Suppl. Fig. S4b, respectively).

Hyperparameters α and β were adjusted via grid search. We aimed at a trade off between reconstruction performance (*SSIM*≥0.6 or *MSE*≤0.01) and regularizations (mitigation of overfitting), which we found for combinations of α= 10^3^, 10^4^ and β = 0–1/16. Next, we performed a permutation test with 10,000 repeats to check if the models trained with images from the upper visual field learned colour-opponent kernels more frequently than those trained with lower visual field images. We compared the number of colour-opponent kernels generated under the two input conditions using a two-sided permutation test.

