## Supplementary Figures for "Mouse retinal specializations reflect knowledge of natural environment statistics"

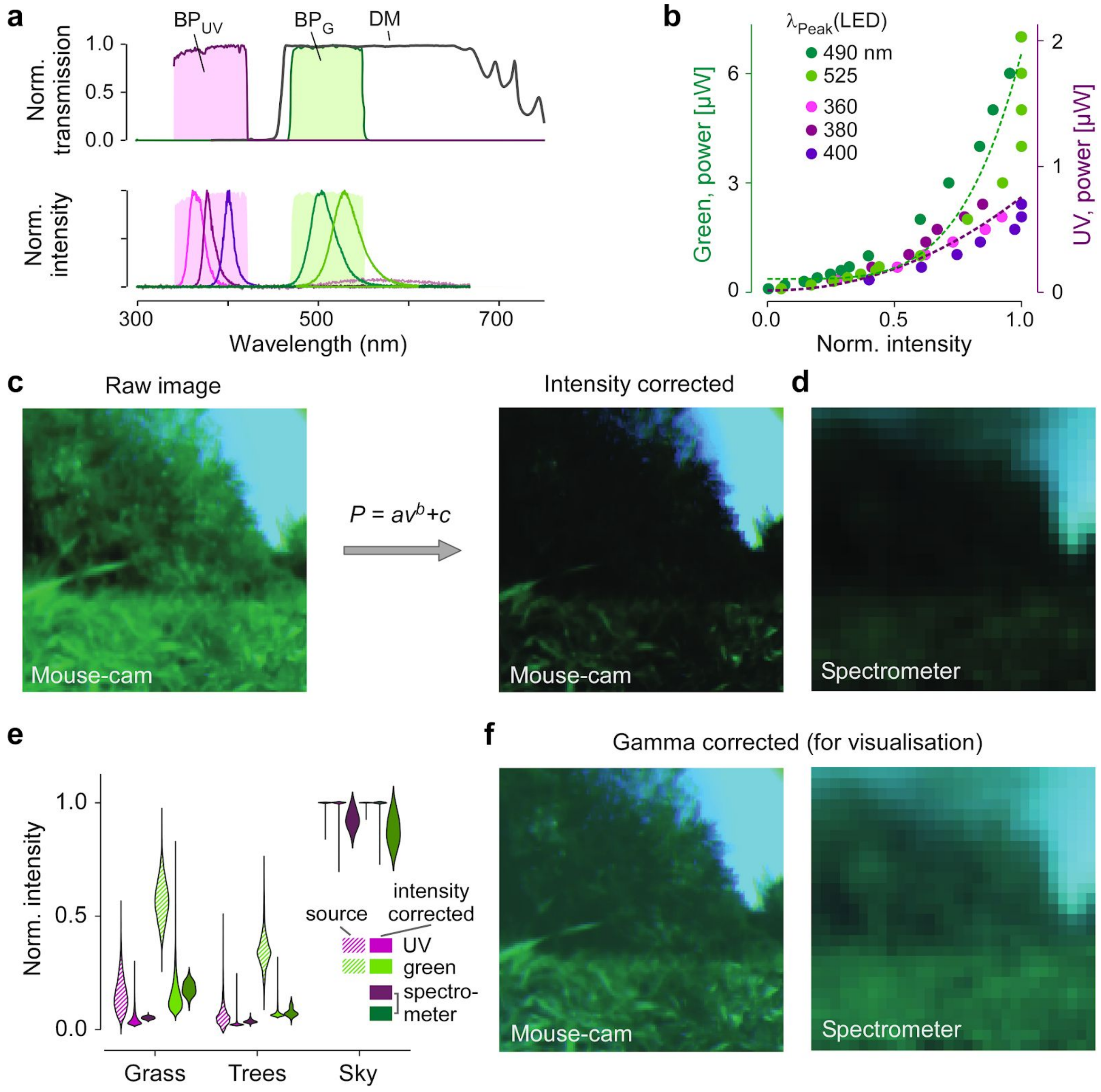


***Supplementary Figure S1 | Intensity correction of the two camera channels****. **a**, Transmission spectra of bandpass filters in UV (BP$_{UV}$) and green (BP$_G$) channels, dichroic mirror (DM; top), and emission spectra of calibration LEDs (bottom; for peak wavelengths, see (b)), **b**, Power of the different LEDs (datapoints; measured with a power meter) as a function of normalised intensity reported by the UV-sensitive and green-sensitive camera (Methods). **c**, Example frame before (left; raw image) and after application of inverse-gamma curves (right; intensity-corrected) from (b). **d**, Same scene recorded using a scanning spectrometer, as described earlier (Baden et al., 2013). **e**, Comparison of intensity distributions (violin plots) of scene "components" (grass, trees, sky) between spectrometer and camera data before and after calibration. Note that due to the limited dynamic range of the camera chip, saturation of pixels in the sky region cannot be completely avoided, hence the narrow distribution of the "Sky" values for the camera measurements. **f**, Images from (c, right) and (d) with gamma correction applied for visualisation (Methods).*

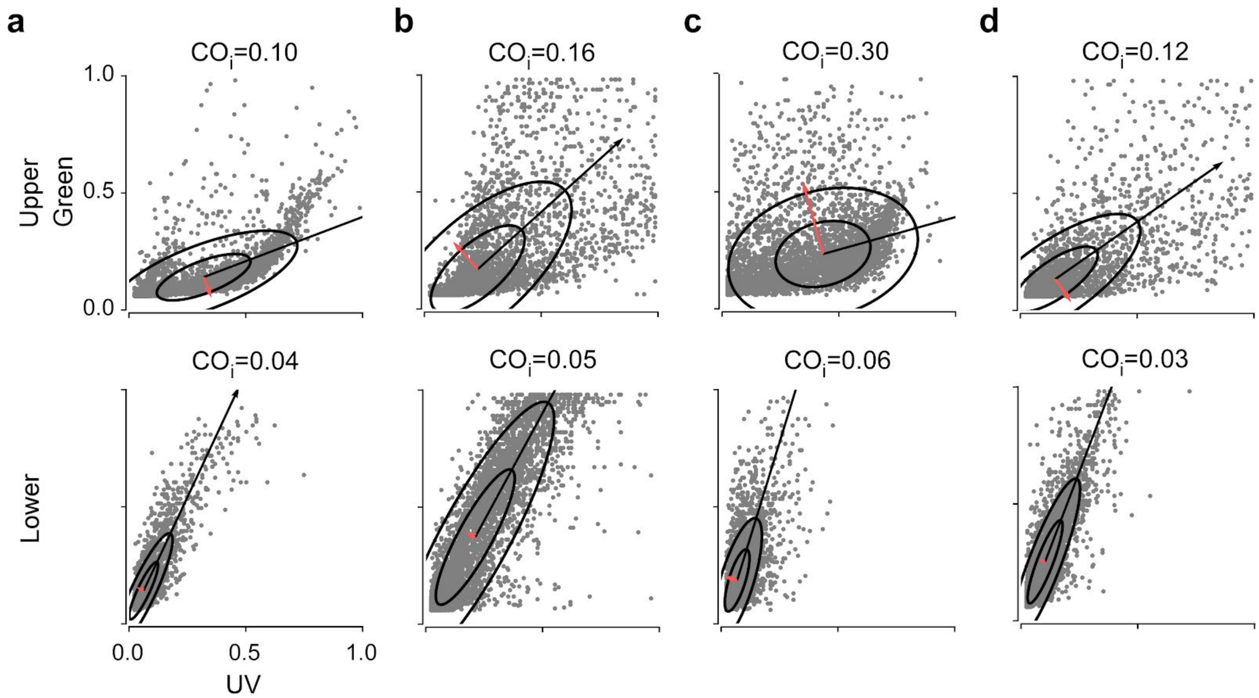


***Supplementary Figure S2 | The colour opponency index (COi) estimated with principal component analysis (PCA) is higher in the upper compared to the lower visual field.*** *PCA is classically used to disentangle achromatic (1st PC) from colour-opponent (2nd and higher PCs) dimensions (Buchsbaum and Gottschalk, 1983).* ***a-d****, 2D plots of 1st and 2nd principle components for intensities of image crops from the upper (top row) and lower (bottom row) visual field. Each dot represents a green-UV intensity pair. Data was fitted with bivariate normal distribution, with ellipses representing 1 and 2 SD. Black and red arrow indicate first and second eigenvector, reflecting the achromatic and the colour-opponent transforms, respectively. For movies, see Fig. 2a and Table 4; (a) top-left, “20190329_13_3”; (b) centre-left, “20180713_13_1”; (c) centre-right, “20180905_14_2”; (d) bottom-left, “20190329 _12_1”). These results are consistent with the idea that chromatic signals are decorrelated at the retinal level (Abbasi-Asl et al., 2016; Graham et al., 2006) and in line with recent studies that suggest a predominance of colour opponent circuits in the ventral mouse retina (Nadal-Nicolás et al., 2020; Szatko et al., 2020)*.

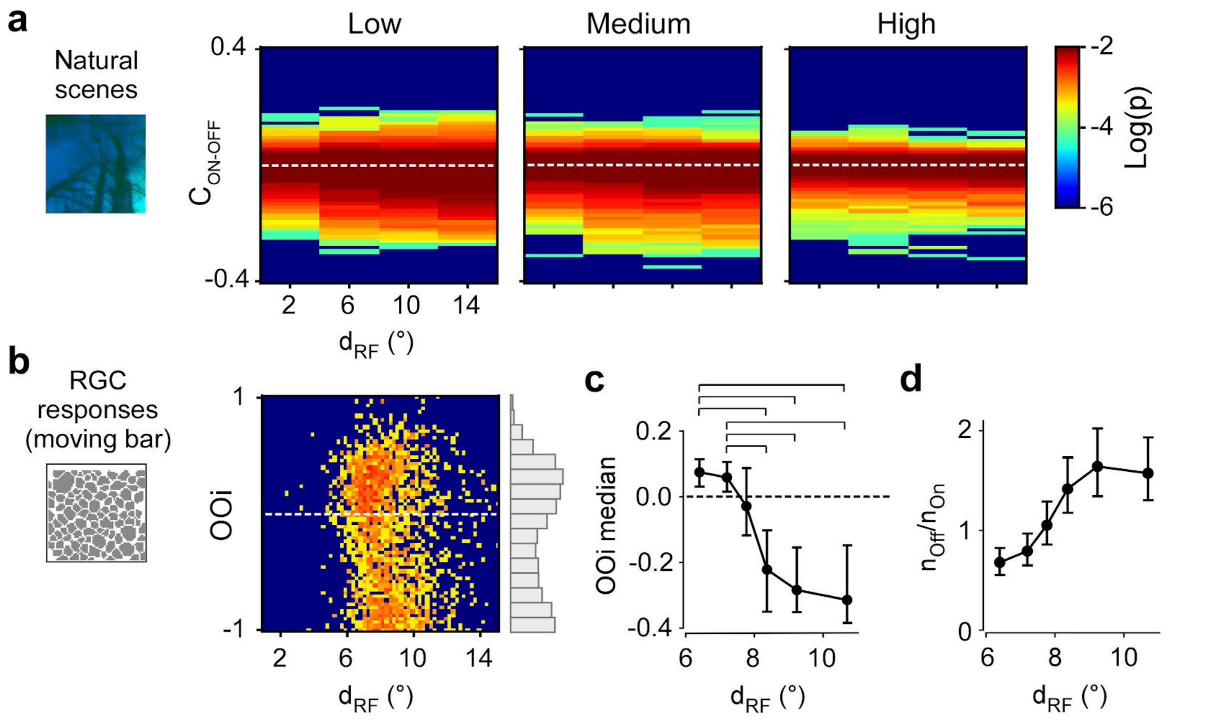


***Supplementary Figure S3 | Bias towards dark contrasts in the upper visual field is reflected in a higher frequency of larger-RF Off RGCs in the ventral retina**. **a**, 2D histogram showing On-Off contrast ($C_{On\text{-}Off}$) measured in upper visual field image crops (same as used for $C_{RMS}$ and $C_{On\text{-}Off}$ in Figs. 3 and 4, respectively) as a function of RF kernel diameter ($d_{RF}$). **b**, 2D histogram showing On-Off index (OOi; see Methods) as a function of $d_{RF}$ for n=2,380 RGCs recorded in the ventral retina (dataset from Baden et al., 2016). Right: Histogram of OOi frequency across RF diameters. **c**, Median OOi as a function $d_{RF}$ (brackets indicate $p<0.0001$; permutation test). **d**, Ratio of Off vs. On RGCs as a function of $d_{RF}$ (with Off and On cells defined as OOi <0 and OOi > 0, respectively). Panels (c,d): Same dataset as in (b); cells divided equally into 6 groups based on their $d_{RF}$; error bars represent 2.5-to-97.5 percentile confidence intervals after bootstrapping.*

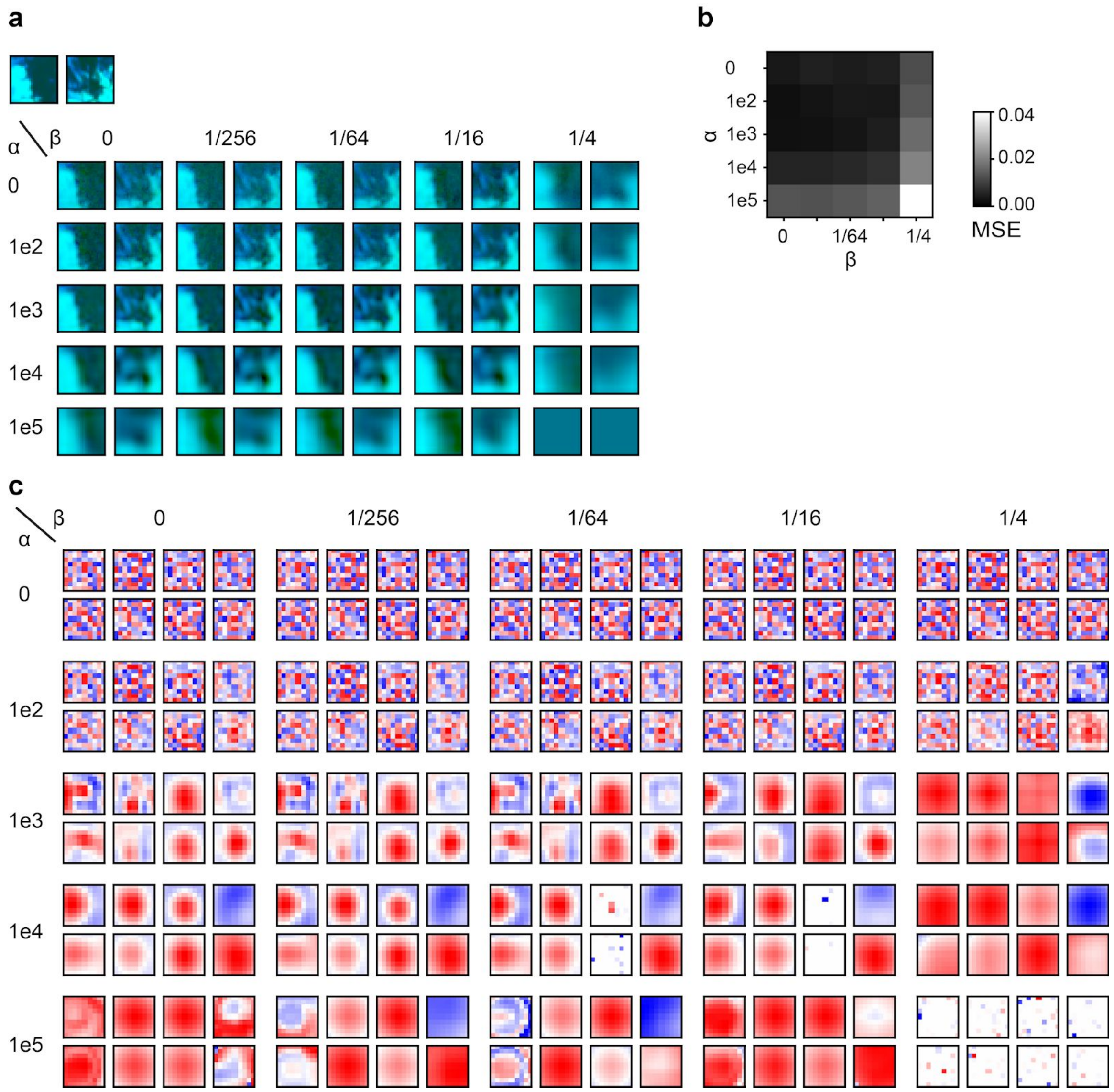


***Supplementary Figure S4 | Training the convolutional autoencoder (CAE) model using image crops from the upper visual field***. ***a***, *Reconstruction of two exemplary image crops (upper left corner) under different regularization values for α (L2) and β (L1).* ***b***, *Reconstruction performance using the mean square error (MSE) as metric. A sudden drop in performance happens for* $\alpha = 10^5$ *or* $\beta = 1/4$*, suggesting over-regularization.* ***c***, *Four of the 16 convolutional kernels under the regularizations from (a). Smooth Gaussian- or Gabor-like kernels are learned for* $\alpha = 10^3$ .. $10^4$ *and* $\beta = 0 \ldots 1/16$*. For details, see Results and Methods.*
